# Correcting dilated cardiomyopathy with fibroblast-targeted p38 deficiency

**DOI:** 10.1101/2023.01.23.523684

**Authors:** Ross C. Bretherton, Isabella M. Reichardt, Kristin A. Zabrecky, Alex J. Goldstein, Logan R.J. Bailey, Darrian Bugg, Timothy S. McMillen, Kristina B. Kooiker, Galina V. Flint, Amy Martinson, Jagdambika Gunaje, Franziska Koser, Elizabeth Plaster, Wolfgang A. Linke, Michael Regnier, Farid Moussavi-Harami, Nathan J. Sniadecki, Cole A. DeForest, Jennifer Davis

## Abstract

Inherited mutations in contractile and structural genes, which decrease cardiomyocyte tension generation, are principal drivers of dilated cardiomyopathy (DCM)– the leading cause of heart failure^1,2^. Progress towards developing precision therapeutics for and defining the underlying determinants of DCM has been cardiomyocyte centric with negligible attention directed towards fibroblasts despite their role in regulating the best predictor of DCM severity, cardiac fibrosis^3,4^. Given that failure to reverse fibrosis is a major limitation of both standard of care and first in class precision therapeutics for DCM, this study examined whether cardiac fibroblast-mediated regulation of the heart’s material properties is essential for the DCM phenotype. Here we report in a mouse model of inherited DCM that prior to the onset of fibrosis and dilated myocardial remodeling both the myocardium and extracellular matrix (ECM) stiffen from switches in titin isoform expression, enhanced collagen fiber alignment, and expansion of the cardiac fibroblast population, which we blocked by genetically suppressing p38α in cardiac fibroblasts. This fibroblast-targeted intervention unexpectedly improved the primary cardiomyocyte defect in contractile function and reversed ECM and dilated myocardial remodeling. Together these findings challenge the long-standing paradigm that ECM remodeling is a secondary complication to inherited defects in cardiomyocyte contractile function and instead demonstrate cardiac fibroblasts are essential contributors to the DCM phenotype, thus suggesting DCM-specific therapeutics will require fibroblast-specific strategies.

## Main

Dilated cardiomyopathy (DCM) is a leading cause of heart failure worldwide that arises from a cadre of insults including inherited mutations in contractile or structural proteins expressed in cardiomyocytes (*1*, *2*, *5*). The clinical hallmarks of DCM are reduced systolic function, thinning of the myocardium, enlargement of the left ventricular chamber, and fibrosis. Despite a robust prevalence of DCM in the population (1 in 250 individuals), there are limited treatment options and, as of yet, no cure (*1*, *6*–*8*). Common to the worst clinical outcomes for DCM is fibrosis, which can precede and exacerbate cardiac structural remodeling (*9*–*11*). When it comes to mitigating fibrosis, first-in-class pharmaceutics for DCM such as myosin modulators have underperformed at correcting fibrotic remodeling in mice and clinical trials, thus tempering their therapeutic value (*5*, *12*). Remodeling of the heart’s extracellular matrix (ECM), such as during the fibrotic response to stress, is primarily regulated by cardiac fibroblasts of the *Tcf21* and *Pdgfra* lineage (*13*–*16*). While cardiac fibroblasts can be activated chemically to secrete fibrotic ECM, these cells are also highly sensitive to mechanical signals including substrate stiffness, alignment, and stretch (*4*). Given that impaired cardiomyocyte force generation is a primary determinant of DCM severity and that fibroblasts are physically coupled to cardiomyocytes via the ECM (*17*), this study examined the hypotheses that fibroblasts function as mechanical rheostats by structurally and biochemically tuning the material properties of the extracellular environment to compensate for disease-linked perturbations to cardiomyocyte force generation and that these adaptations are essential second drivers of the DCM phenotype.

### Adaptive structural reorganization and stiffening of the ECM precedes myocyte dilation

To determine whether the material properties of the ECM adapt in response to DCM-linked reductions in cardiomyocyte force generation, a previously reported I61Q mutant cardiac troponin C (cTnC) transgene was specifically expressed in cardiomyocytes using a doxycycline-repressible alpha-myosin heavy chain promoter (αMHC, **Fig. 1a**)(*2*). The I61Q point mutation lowers the binding affinity of Ca^2+^ to cTnC and thereby desensitizes myofilaments to Ca^2+^, reducing cardiomyocyte force production on a beat-to-beat basis (*18*, *19*). This change is apparent by echocardiography, which showed reduced ejection fraction in the hearts of I61Q transgenic mice by 2 months of age relative to the wild type (WT) control group, which consisted of non-transgenic and tetracycline transactivator (tTA) transgenic littermates that were previously shown to be statistically equivalent (**Fig. 1b**) (*2*). Dilated structural remodeling in the hearts of I61Q transgenic mice was first detected at 4 months of age, as indicated by increased diastolic left ventricular (LV) chamber dimensions and heart-to-body weight ratios (**Fig. 1c-d**).This dilated phenotype was also observed in cardioplegia-arrested myocardial sections at the same timepoint (**Fig. 1e**). Concomitant with structural dilation, hearts from I61Q transgenic mice had mild interstitial fibrosis that was histologically undetectable early in the disease process but emerged by 4 months of age (**Fig. 1f-g**). As the biochemical composition and density of the ECM determines its mechanical properties (*20*), label-free data-independent acquisition (DIA) mass spectrometry of decellularized cardiac ECM from I61Q cTnC transgenic and WT mice was performed at the onset of dilation (**Fig. S1**). The relative abundance of primary ECM constituents was largely unchanged between genotypes (**Fig. S1a**). However, closer examination revealed 22 core matrisome proteins were differentially expressed in the cardiac ECM of WT and I61Q transgenic mice, including several laminin subtypes and type VI collagen, an established biomarker of heart failure (**Fig. S1b**) (*21*–*23*).

**Fig. 1.**
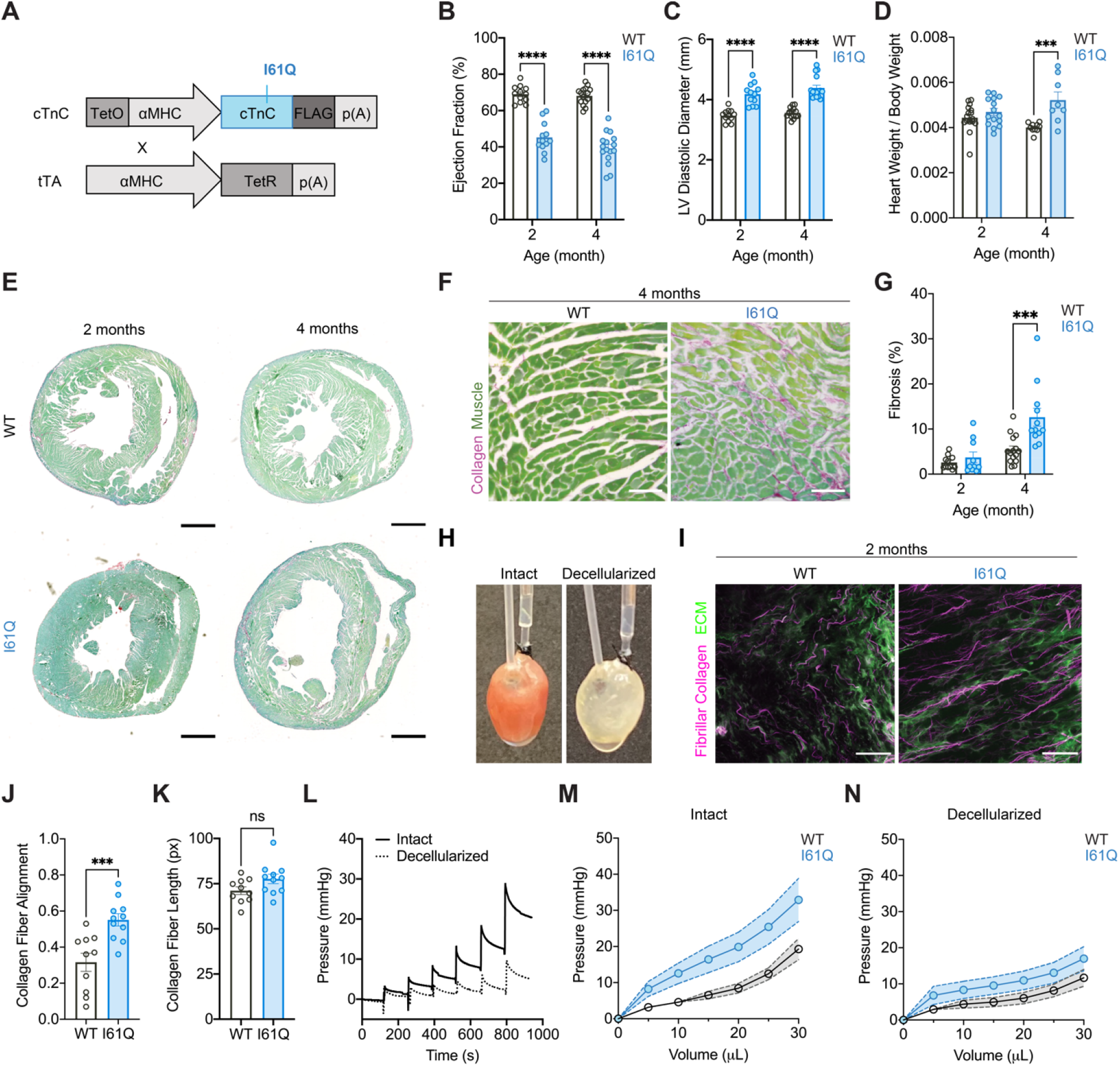
Reduced myocyte tension generation aligns collagen and stiffens the myocardium prior to overt fibrosis and dilated structural remodeling. **(A)** Schematic of the genetic crosses used to generate I61Q cTnC transgenic mice. Mice with cardiomyocyte specific expression of FLAG-tagged I61Q mutant cardiac troponin c (cTnC) transgene driven by a tetracycline regulated α-myosin heavy chain (αMHC) promoter were crossed with tetracycline transactivator (tTA) mice which causes constitutive expression of the I61Q mutant cTnC transgene. **(B)** Quantification of left ventricular ejection fraction and **(C)** diastolic chamber diameter by echocardiography at 2 (I61Q n=12, WT n=12) and 4 (I61Q n=16, WT n=15) months of age. **(D)** Quantification of hypertrophy by heart weight to body weight ratio of I61Q mice (2 month n=15, 4 month n=8) and WT controls (2 month n=17, 4 month n=10). **(E)** Representative images of cardiac paraffin sections stained with picrosirius red-fast green (PSR/FG, scale bar=1mm) at 2 and 4 months of age. **(F)** Representative 20x images (scale bar=50μm) at 4 months and **(G)** quantification of PSR/FG staining on myocardial sections from 2 (WT n=11, I61Q n=10) and 4 months (WT n=15, I61Q n=12). **(H)** Representative images of an intact (left) and decellularized (right) heart in a modified Langendorff working heart preparation for passive mechanical measurements. **(I)** Representative two-photon images of decellularized hearts (scale bar=100μm) showing fibrillar collagen (magenta) and ECM autofluorescence (green). **(J)** CurveAlign quantification of collagen fiber alignment and **(K)** length from 2-month-old I61Q (n=10) hearts and WT controls (n=11). **(L)** Representative developed pressure traces from stepwise inflation of a balloon inside a blebbistatin-treated intact (dark line) and decellularized (straight line) heart. **(M)** Pressure volume curves of intact and **(N)** decellularized mouse hearts at 2 months (n=7 both genotypes). Data are mean ± SEM, ns=not significant, ***p<0.005, ****p<0.001 by 2-way ANOVA with Holm-Sidak’s multiple comparisons test (**B-D, G**) or two-tailed unpaired t-test (**J,K**).

Structural reorganization of collagen fibers is another mechanism for adaptive mechanical tuning of the ECM (*24*). Hence, label-free second harmonic generation (SHG) microscopy was used to evaluate collagen structure and organization in whole mount decellularized I61Q transgenics and WT hearts (**Fig. 1h**). While differences in fibrosis by conventional histology were unresolved at 2 months of age, SHG imaging exposed robust increases in circumferential collagen fiber alignment in I61Q transgenic hearts at this early timepoint (**Fig. 1i-j**), suggesting structural reorganization of collagen fiber topography is a proximal compensatory response to reduced force generation by cardiomyocytes. Collagen fiber length, while longer on average in I61Q transgenics, was not significantly different from WT controls (**Fig. 1k**), indicating the observed topographical changes were not simply a product of collagen elongation. Transitioning to a highly aligned collagen fiber topography typically increases both the anisotropic strength of the ECM and force transmission of myocardial tissue, which in turn increases stiffness and preserves the heart’s ability to contract (*25*). Such an adaptation would be a vital compensatory response to the I61Q cTnC-dependent loss in cardiomyocyte force generation.

To determine if the myocardium and cardiac ECM stiffen in response to I61Q cTnC expression, the passive mechanical properties of intact and decellularized hearts from the same experimental animal were measured using a modified Langendorff assay. Here, intact hearts from I61Q transgenic and WT mice were subjected to retrograde perfusion with Krebs-Henseleit buffer containing the myosin inhibitor blebbistatin to negate any stiffness from attached cross-bridges, and then a balloon was inserted into the left ventricle for volumetric inflation of the chamber in a stepwise manner. The balloon pressure was recorded during each inflation step, which exhibited a maximal pressure required to achieve the initial volume change and followed by a gradual decrease in pressure due to viscoelastic relaxation (**Fig. 1l**). Following the intact measurements, hearts were decellularized and the assay repeated to measure passive mechanical properties of the ECM in the same preparation. Both intact and decellularized preparations from I61Q mice required higher maximal pressures per inflation, indicating both the myocardium and ECM are stiffer relative to WT (**Fig. 1m-n, Fig. S2**). As they play a major regulatory role in dictating myocyte stiffness, titin isoforms and post-translational modifications were surveyed by Western blot analysis. Here, the stiffer N2B isoform was significantly upregulated in I61Q hearts whereas phosphorylation of the serine residue at position 267 (S267) in the N2B unique sequence, which enhances myocyte compliance, was reduced (**Fig. S3**) (*26*, *27*). Collectively, these results demonstrate that architectural reorganization of collagen fibers and myocardial stiffening precede dilated remodeling and fibrosis associated with DCM.

### Cardiac fibroblasts compensate for reduced cardiomyocyte force generation through proliferation rather than activation

While titin composition was statistically altered in I61Q cardiomyocytes, the modest effect size prompted deeper examination of the basis for structural realignment and stiffening of the myocardium and ECM in I61Q transgenic hearts. A potent determinant of ECM stiffness is fibroblast activation and conversion to a myofibroblast state, which is an essential cellular process underlying fibrosis (*28*). To determine whether mutant I61Q cTnC induces fibroblasts to activate and transition to a profibrotic myofibroblast state, cardiac fibroblasts were isolated from I61Q transgenic and WT hearts for primary culture, stimulated with recombinant TGFβ1(*4*), and the percentage of the population that had smooth muscle α-actin (αSMA)-positive stress fibers quantified. This assay revealed no differences between genotypes at baseline or in response to TGFβ1, suggesting fibroblasts from I61Q transgenic hearts have not differentiated into myofibroblasts, nor were they sensitized to activation signals (**Fig. S4a-b**). Since cell behaviors *in vitro* are often not recapitulated *in vivo*, myocardial sections from I61Q cTnC and WT mice were examined for the presence of activated myofibroblasts by quantifying the number of cells that were positive for two discriminating myofibroblast markers: αSMA and platelet-derived growth factor alpha (PDGFRα). Surprisingly, no significant fibroblast-to-myofibroblast conversion was evident even after the onset of fibrosis in I61Q hearts (**Fig. 2a-b**). *Acta2* gene transcription was assayed in purified cardiac fibroblasts, but again no significant differences were observed between genotypes, further suggesting that cardiac fibroblasts were not transitioning to myofibroblasts (**Fig. 2c**).A dual color fluorescent reporter was also used to trace activated fibroblasts *in vivo* with a *Postn* Cre-driver (*Postn^iCre^-mT/mG*) (*13*, *29*), which was able to detect pockets of activated *Postn^+^* cells in I61Q transgenic hearts, albeit at 8 months of age which is long after the I61Q cTnC hearts dilate and turn towards decompensation (**Fig. 2d-e, Fig. S4c**) (*2*). These results suggest that (1) the canonical fibrotic process of fibroblast activation and myofibroblast formation only occurs in this inherited DCM model once the heart progresses to failure, and (2) fibroblast activation to an intermediate or fully matured αSMA^+^ myofibroblast state is not essential for structural realignment and stiffening of the ECM.

**Fig. 2.**
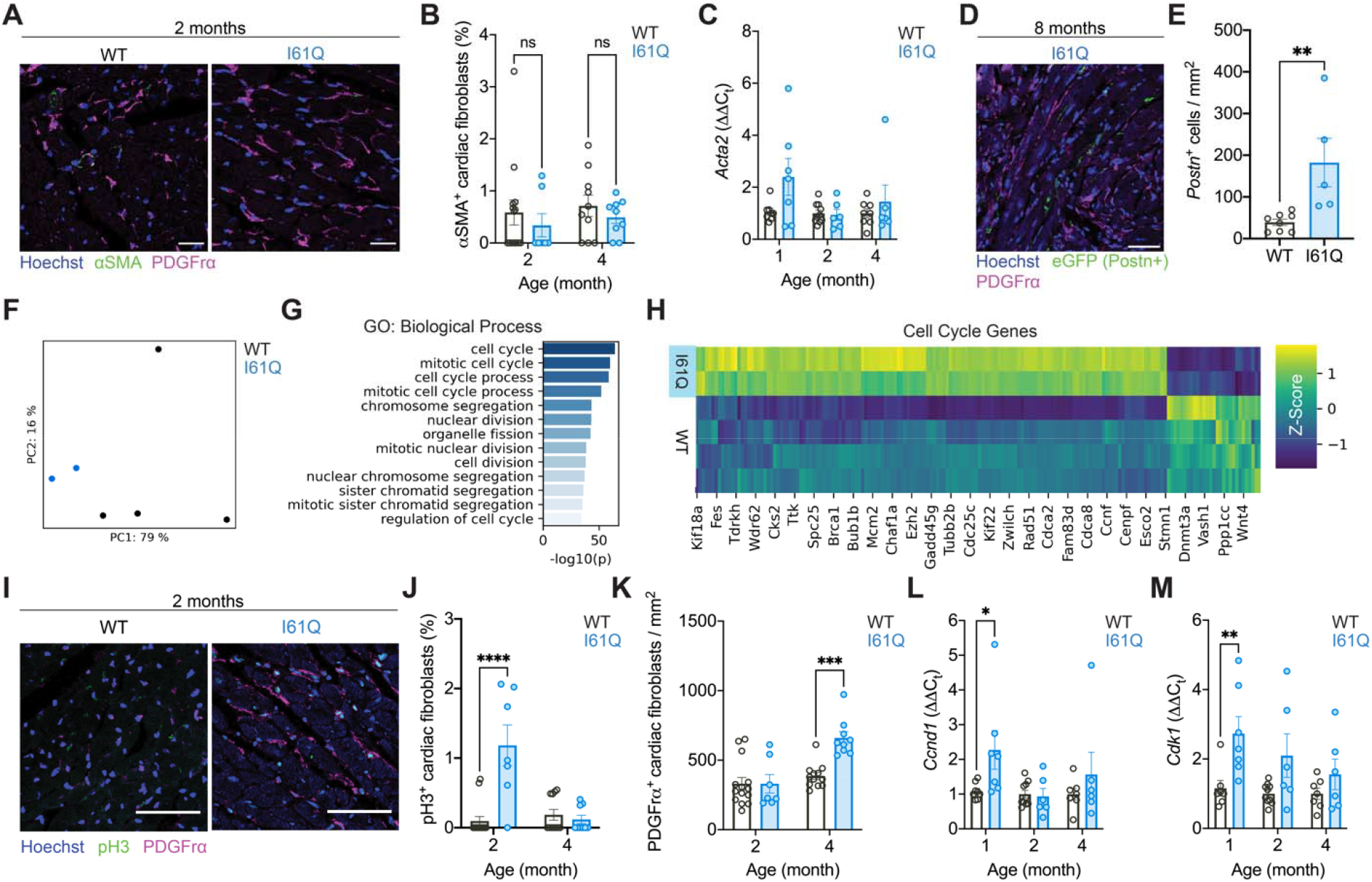
DCM-linked I61Q mutant cTnC initiates cardiac fibroblast proliferation rather than activation. **(A)** Representative images of 2-month-old WT and I61Q cardiac sections stained for smooth muscle α-actin (αSMA) and platelet-derived growth factor receptor α (PDGFRα). **(B)** Quantification of immunofluorescent imaging for the percentage of fibroblasts (PDGFRα^+^) expressing αSMA (2 month n=14 WT, n=7 I61Q, 4 month n=10 WT, n=9 I61Q). **(C)** RT-PCR for Acta2 transcript in magnetically sorted cardiac fibroblasts from WT (n=7,10,8) and I61Q (n=7,6,6) hearts at 1, 2, and 4 months of age. **(D)** Representative image and **(E)** quantification of Postn^+^ cell density in I61Q (n=5) and WT (n=8) hearts at 8 months of age. **(F)** Principal component analysis of cardiac fibroblast transcriptomes from 4-month-old wild type (WT, n=4) and I61Q (n=2) mice. **(G)** Significant gene ontology biological processes (GO:BP) enriched in differentially regulated genes from I61Q fibroblasts. **(H)** Heatmap showing expression levels of cell cycle genes. **(I)** Representative images from immunofluorescent staining for phospho-histone H3 (pH3) and platelet-derived growth factor receptor alpha (PDGFRα) in I61Q and WT myocardium. **(J)** Quantification of fibroblast pH3 positivity rates and **(K)** density (n same as **A**). **(L)** RT-PCR for the cell cycle transcripts *Ccnd1* and **(M)***Cdk1* on magnetically sorted fibroblast samples isolated from WT and I61Q hearts (n same as **C**). Data are mean ± SEM, ns=not significant, *p<0.05,**p<0.01,***p<0.005,****p<0.001 by 2-way ANOVA with Holm-Sidak’s multiple comparisons test (**B,C,J-M**) or two-tailed unpaired t-test (**E**). All scale bars (**A,D,J**) = 50μm.

ECM remodeling could also be driven by fibroblast proliferation or other functional state changes (*30*–*33*). Hence, transcriptome profiling by RNA sequencing (RNAseq) was performed on cardiac fibroblasts isolated from 4-month-old I61Q mice and WT controls. Though the fibroblast is not genetically manipulated in the I61Q mice, principal component analysis (PCA) separated the animal genotypes from which fibroblasts were derived on the first principal component (PCA1) and accounted for 79% of sample variance (**Fig. 2f**). Differential gene expression analysis identified 363 significantly upregulated genes and 449 significantly downregulated genes in I61Q fibroblasts relative to controls (**Data S1**). Pathway enrichment analysis with g:Profiler found that all ten of the top enriched pathways were related to cell cycle (**Fig. 2g**) (*34*). Of the genes within this category, a variety of critical cell cycle regulators were upregulated in I61Q fibroblasts, including several cyclin genes (*Ccnb1*, *Ccnb2*, *Ccnd1*, *Ccne2*, *Ccnf*), cyclin dependent kinase 1 (*Cdk1*), marker of proliferation Ki-67 (*Mki67*), and aurora kinase (*Aurka*) (**Fig. 2h**). To assess whether altered levels of cell cycle markers were driving heightened proliferation in I61Q fibroblasts, cardiac sections were stained for the fibroblast marker PDGFRα and cell cycle marker phospho-histone H3 (pH3), which demonstrated that fibroblasts in I61Q hearts had heightened proliferation signals at the 2-month timepoint and a *bona fide* increase in PDGFRα^+^ fibroblast density by 4 months of age (**Fig. 2i-k**). Significant upregulation of *Ccnd1* and *Cdk1* transcripts was captured even earlier in purified fibroblast preparations from 1-month-old mice (**Fig. 2l-m**).

Biochemical cues present in the cardiac ECM can also modulate cell proliferation, suggesting that I61Q ECM could further drive fibroblast proliferation in positive feedback (*35*). To test this, cardiac fibroblasts were encapsulated in poly(ethylene glycol) (PEG)-based hydrogels modified to present biochemical ECM cues to cells in an environment with conserved mechanics. To achieve this, pepsin-digested ECM from WT or I61Q hearts was functionalized with 4-azidobutyric acid N-hydroxysuccinimide ester and covalently decorated onto a step-growth PEG hydrogel by cytocompatible copper-free click chemistry (*36*, *37*). I61Q fibroblasts cultured within these soft hydrogels (~2 kPa storage modulus) no longer retained their hyperproliferative phenotype, but instead fibroblast proliferation was modulated by the chemical constituents of the ECM (**Fig. S5a**).In a screen of 36 combinations of ECM proteins in array on a soft (~10 kPa) polyacrylamide gel, type VI collagen supported more efficient cardiac fibroblast adhesion to the substrate and did so in synergy with laminin (**Fig. S5b-e**). These two proteins were enriched in our I61Q ECM proteomics, underscoring a potential role for matrix signals in initiating fibroblast proliferation and advancing the DCM phenotype.

### Fibroblast proliferation is sufficient to drive collagen compaction and tissue alignment

Adaptive alignment of fibrillar collagen and ECM stiffening in inherited DCM could result from traction forces exerted by cells on the matrix as the myocardium becomes progressively volume overloaded(*38*). Indeed, hyperproliferative I61Q fibroblasts compacted their surrounding ECM to a greater extent than those from WT hearts following encapsulation in free-floating collagen gels (**Fig. 3a-b**). To test whether proliferation was essential to gel compaction, a small-molecule cyclin-dependent kinase inhibitor (CDKi) dinaciclib was delivered in the culture media. CDKi treatment reduced proliferation of cardiac fibroblasts from both genotypes to similarly low levels (**Fig. 3c**) and blocked genotype-dependent gel compaction (**Fig. 3d**), demonstrating that increased fibroblast numbers rather than greater contractile function of the cell caused the compaction. To further confirm that gel compaction was due to proliferation rather than ECM degradation, a set of cell-laden collagen gels were also treated with marimastat, a broad-spectrum inhibitor of matrix metalloproteinases, which had no significant effect on I61Q fibroblast proliferation or gel compaction for either fibroblast genotype (**Fig. S6**). To examine the effects of load on tissue alignment and stiffness, fibroblasts were seeded into fibrin gels suspended between a flexible and a rigid post made of polydimethylsiloxane (PDMS) (**Fig. 3e**). Similar to the collagen gels, fibrin tissues seeded with cardiac fibroblasts from I61Q transgenic hearts had increased compaction and generated more passive tension, as measured by the magnitude of PDMS post deflection (**Fig. 3f-g**).Concomitant with the heightened passive tension, I61Q fibroblasts were more aligned within the tissues, suggesting that fibroblast proliferation causes tissue compaction, which thereby promotes cellular alignment (**Fig. 3h**).

**Fig. 3.**
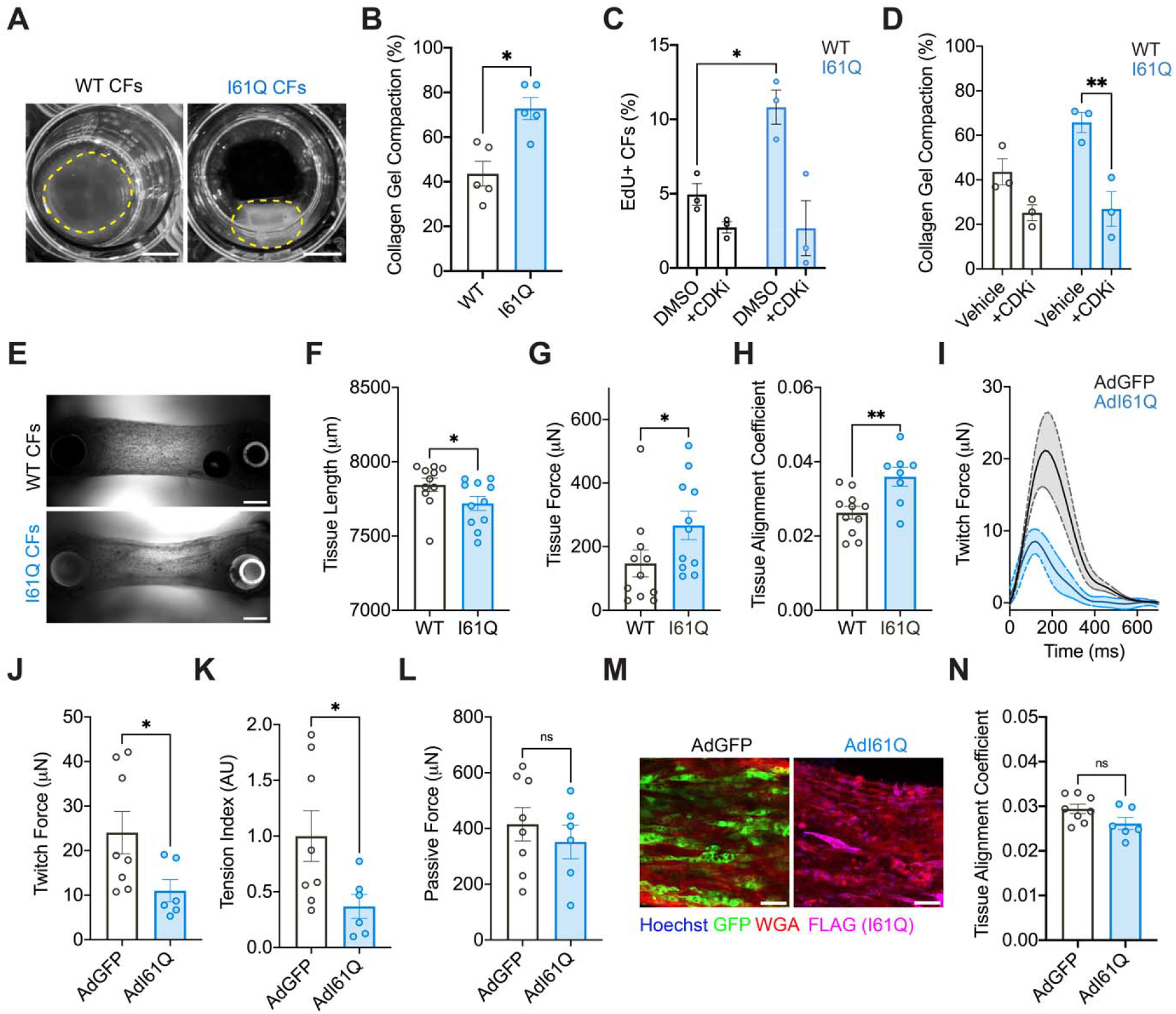
Cardiac fibroblasts derived from I61Q cTnC transgenic hearts aligns engineered cardiac tissues. **(A)** Representative images (scale bar=5 mm) and **(B)** quantification of the compaction of free-floating collagen gels seeded with cardiac fibroblasts (CFs) isolated from I61Q cTnC transgenic and WT hearts (n=5 per genotype). **(C)** Quantification of cardiac fibroblast proliferation by EdU incorporation in the genotypes indicated. **(D)** Quantification of the compaction of free-floating collagen gels seeded with cardiac fibroblasts isolated from I61Q cTnC transgenic and WT hearts plus administration of dinaciclib (CDKi) or vehicle (DMSO) in the culture media (n=3 per genotype). **(E)** Representative images of cardiac fibroblast-seeded fibrin tissues mounted between PDMS posts (scale bar=1 mm). **(F)** Quantification of length and **(G)** force production by tissues seeded with cardiac fibroblasts from I61Q cTnC transgenic or WT mice (n=11 per genotype). **(H)** Quantification of cellular and ECM alignment in fibrin tissues seeded cardiac fibroblasts derived from I61Q cTnC transgenic and WT hearts by wheat germ staining (n=11 WT, n=8 I61Q). **(I)** Average twitch forces generated by engineered heart tissues (EHTs) 2 weeks after cardiomyocytes were adenovirally transduced with either control (AdGFP, n=8) or I61Q mutant cTnC (AdI61Q, n=6). **(J)** Quantification of twitch force, **(K)** tension index, and **(L)** passive force generation by EHTs. **(M)** Representative images (scale 50μm) and **(N)** quantification of EHT alignment by wheat germ staining. Data are mean ± SEM, ns=not significant, *p<0.05,**p<0.01 by 2-way ANOVA with Holm-Sidak’s multiple comparisons test (**C,D**) or two-tailed unpaired t-test (**B, F-H, J-L, N**).

Though traction forces from fibroblast could contribute to ECM alignment, early stiffening of cardiomyocytes and altered hemodynamic loading during DCM pathogenesis could also produce the traction needed to align and lengthen collagen fibers. To study the effects of the mutant I61Q cTnC on tissue alignment in the absence of hemodynamic load, naïve neonatal rat cardiomyocytes were seeded into engineered heart tissues (EHTs) between PDMS posts and adenovirally transduced with FLAG-tagged I61Q cTnC (AdI61Q) or green fluorescent protein (AdGFP) as a transduction control. Transduced EHTs were cultured for two weeks prior to analyzing contractile output, passive tension generation, and tissue alignment. Here, AdI61Q EHTs functionally phenocopied the I61Q transgenic mice (*2*), including reduced twitch force and reduced area under the twitch curve, previously referred to as the tension index (**Fig. 3i-k**). Notably absent from the EHT phenotype was any difference in passive tension generation (**Fig. 3l**), in contrast to what was observed in tissues engineered with fibroblasts from I61Q hearts (**Fig. 3g**).Tissue alignment was similarly unaffected by cardiomyocytes transduced with AdI61Q (**Fig. 3m-n**).Taken together these experiments demonstrate that in the absence of fibroblast-generated passive tension or hemodynamic loading, I61Q expression by the cardiomyocyte alone is insufficient to produce myocardial tissue alignment and stiffness.

### DCM is reversed by targeted deletion of p38 in cardiac fibroblasts

Based on the finding that fibroblast-dependent ECM alignment and tissue stiffening precede fibrosis, it was hypothesized that therapeutic interventions for DCM should disrupt fibroblast mechanotransduction and function. Yet known was how cardiac fibroblasts sense and transduce the effects of I61Q mutant cTnC on myocyte function. It was observed that the focal adhesions were significantly larger and more elongated in cardiac fibroblasts isolated from I61Q transgenic hearts, which phenocopies naïve fibroblasts cultured on engineered biomimetics of aligned collagen topography (**Fig. S7a-c**) (*39*). This suggests that the fibroblasts may sense I61Q cTnC-dependent perturbations to the mechanical environment via ECM and integrin signaling. Previous findings from our lab demonstrated that extracellular signals governing fibroblast function are transduced by p38 mitogen-activated protein kinase (p38 MAPK) signaling (*39*–*41*). Hence, cardiac fibroblast-specific p38 activity was examined in I61Q transgenic mice at the 2-month timepoint when fibroblasts are proliferative and adaptive structural alignment and stiffening occurs. By Western blot analysis, both total and phosphorylated p38 levels were upregulated in purified fibroblasts from I61Q hearts when compared to WT (**Fig. 4a**). Moreover, nuclear translocation of p38 was greater in cultured fibroblasts from I61Q transgenic hearts relative to WT (**Fig. S7d-e**) providing further evidence that the I61Q cTnC transgene enhances p38 activity in cardiac fibroblasts.

**Fig. 4.**
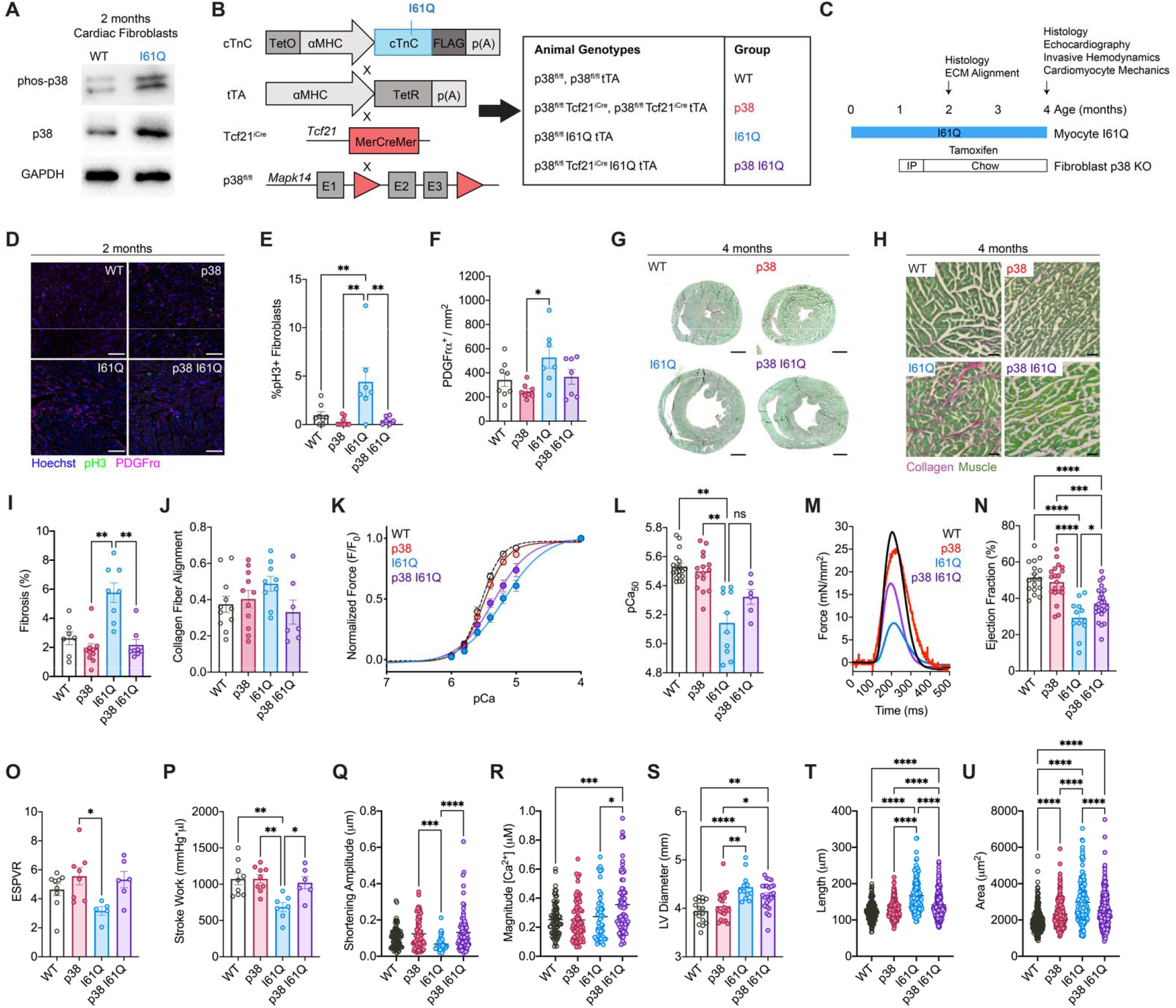
Fibroblast-specific deletion of p38 corrects cardiac dilation and systolic dysfunction in I61Q cTnC transgenic mice. **(A)** Quantification of p38 abundance and phosphorylation by Western blot on lysates from purified cardiac fibroblasts isolated from I61Q cTnC transgenic and WT hearts. **(B)** Schematic showing the generation of I61Q cTnC transgenic mice with tamoxifen inducible fibroblast-specific p38 deletion and the experimental genotypes derived from the described breeding scheme. Here, I61Q cTnC and tTA transgene were bred with a mouse line containing conditional p38α loss of function (p38^fl/fl^) and a tamoxifen-inducible Cre recombinase knocked into the *Tcf21* locus (Tcf21^iCre^). **(C)** Experimental design schematic showing the I61Q mutant cTnC is expressed just after birth (~ postnatal day 2), mice were allowed to develop normally for 1 month, and then tamoxifen was administered to induce fibroblast-specific p38 excision. Experimental endpoints were at 2- and 4-months of age. **(D)** Representative immunofluorescent staining (scale bar=50μm) for pH3 and PDGFRα in 2-month-old sections, and quantification of **(E)** fibroblast proliferation rates and **(F)** fibroblast density. (WT n=8, p38 n=8, I61Q n=7, p38 I61Q n=7) **(G)** Representative cardiac cross-sections from 4-month-old mice stained with PSR/FG. **(H)** Representative 20x regions of interest (ROIs) and **(I)** quantification of PSR/FG staining (WT n=7, p38 n=11, I61Q n=8, p38 I61Q n=8). **(J)** Quantification of collagen fiber alignment from decellularized WT (n=11), p38 (n=11), I61Q (n=9), and p38 I61Q (n=7) hearts. **(K)** Representative relationship between normalized tension and Ca^2+^ concentration (pCa) and **(L)** Ca^2+^ sensitivity of tension generation (pCa_50_) in membrane permeabilized trabeculae of WT (WT, n=19), p38^fl/fl^-Tcf21^iCre^ (p38, n=15), I61Q cTnC-tTA (I61Q, n=10), and p38^fl/fl^-Tcf21^iCre^-I61Q-tTA (p38 I61Q, n=6) mice. **(M)** Mean twitch forces from intact trabeculae of 4-month-old WT (n=4), p38 (n=3), I61Q (n=4), and p38 I61Q (n=4) mice. **(N)** Quantification of left ventricular ejection fraction measured by echocardiography from WT (n=17), p38 (n=20), I61Q (n=12), and p38 I61Q (n=23) mice. **(O)** Measurement of the end systolic pressure volume relationship (ESPVR) and **(P)** stroke work by invasive hemodynamics from 4-month-old mice (WT n=9, p38 n=9, I61Q n=7, p38 I61Q n=6). **(Q)** Quantification of unloaded sarcomere shortening amplitude (WT n=85, p38 n=87, I61Q n=45, p38 I61Q n=85 cardiomyocytes) and **(R)** Ca^2+^ transient amplitude (WT n=75, p38 n=81, I61Q n=54, p38 I61Q n=66) in isolated intact cardiomyocytes from the described genotypes. **(S)** Quantification of left ventricular diastolic diameter at 4 months of age by echocardiography (n same as **N**). **(T)** Quantification of isolated cardiomyocyte length and **(U)** area from the described genotypes (WT n=250, p38 n=250, I61Q n=199, p38 I61Q n=200). Data are mean ± SEM, ns=not significant, *p<0.05,**p<0.01,,***p<0.005,****p<0.001 by 2-way ANOVA with Holm-Sidak’s multiple comparisons test.

To directly determine if p38-dependent cardiac fibroblast function plays a role in non-ischemic DCM remodeling, I61Q cTnC transgenic mice were crossed with a mouse line that has tamoxifen-inducible loss of p38 function specifically in cardiac fibroblasts, giving rise to four experimental genotypes: WT controls (p38^fl/fl^ or *Tcf21^iCre^*), fibroblast-specific p38 knockouts (p38^fl/fl^-Tcf21^*iCre*^), I61Q cTnC (I61Q-p38^*fl/fl*^), and I61Q cTnC with fibroblast-specific p38 deletion (I61Q-p38^fl/fl^-Tcf21^*iCre*^) (**Fig. 4b**). At weaning, mice from this cross received one week of tamoxifen intraperitoneal injections followed by 10 weeks of tamoxifen chow (**Fig. 4c**), which we have previously shown elicits ~85% recombination efficiency and nearly complete p38 deletion within 2 weeks of tamoxifen induction in cardiac fibroblasts homozygous for the conditional p38 allele and heterozygous for the Tcf21^*iCre*^ knock-in allele(*41*). 2-month-old myocardial sections immunostained with PDGFRα and pH3 antibodies demonstrated that I61Q transgenic mice with cardiac fibroblast specific p38 deletion (I61Q cTnC-p38^fl/fl^-Tcf21^*iCre*^) had a significant reduction in the number of actively proliferating cardiac fibroblasts as demonstrated by the reduction in fibroblasts that were double positive for PDGFRα and pH3 (**Fig. 4d-e**). This loss in cell cycle activity likely underlies the reduction in PDGFRα^+^ fibroblasts per area of the heart observed in most I61Q cTnC-p38^fl/fl^-Tcf21^*iCre*^ hearts (**Fig. 4f**). 4-month-old myocardial cross-sections stained with picrosirius red-fast green also showed that fibroblast-specific loss of p38 function in I61Q transgenic mice (I61Q cTnC-p38^fl/fl^-Tcf21^*iCre*^) corrects ventricular chamber dimensions as well as interstitial fibrosis at the later timepoint (**Fig. 4g-i**). Analysis of collagen fiber alignment by SHG imaging of decellularized hearts from these mice also demonstrated that in most of the I61Q cTnC-p38^fl/fl^-Tcf21^*iCre*^ cohort collagen fibers were on average less aligned like WT controls, although this metric was not yet statistically significant (**Fig. 4j**).

To confirm that alterations in cardiac fibroblast proliferation and matrix phenotype were due to p38-dependent changes in fibroblast function rather than an alteration in the primary myocyte contractile defect incurred from replacing native cTnC with the I61Q mutant, Ca^2+^-activated force generation was measured in demembranated trabecula from all of the experimental genotypes generated from crossing I61Q cTnC transgenic mice with fibroblast-specific p38 knockouts (p38^fl/fl^-Tcf21^*iCre*^). As represented by a marked rightward shift in the isometric cardiac muscle force–Ca^2+^ relationship (**Fig. 4k**), there was an I61Q cTnC transgene-dependent decrease in force generation at half-maximal Ca^2+^ concentrations (pCa_50_) that was retained in I61Q cTnC-p38^fl/fl^-Tcf21^*iCre*^ cardiac muscle when compared to WT and fibroblast-specific p38 knockout (p38^fl/fl^-Tcf21^*iCre*^) controls (**Fig. 4l**). These data demonstrate that expression of I61Q mutant cTnC retains its primary functional defect of desensitizing the myofilaments to Ca^2+^ despite the fibroblast-specific deletion of p38 in I61Q cTnC-p38^fl/fl^-Tcf21^*iCre*^ mice. Twitch forces were also measured in intact cardiac muscle from these mice. Unexpectedly, loss of p38 function in cardiac fibroblasts significantly corrected the I61Q cTnC-dependent impairment of myocyte twitch function in intact I61Q cTnC-p38^fl/fl^-Tcf21^*iCre*^ cardiac muscle preparations, as shown by enhanced force generation throughout the contraction and relaxation phase of the twitch in comparison to the group expressing I61Q cTnC alone (**Fig. 4m**). Functional rescue of the I61Q cTnC phenotype was also seen at the whole heart level by echocardiography in which I61Q cTnC-p38^fl/fl^-Tcf21^*iCre*^ mice had a significant recovery in ejection fraction (**Fig. 4n**). Invasive hemodynamics further confirmed that a p38-dependent modulation of fibroblast phenotype corrects systolic function in I61Q transgenic mice, as end systolic pressure-volume relationship (ESPVR) and cardiac stroke work were fully restored to WT values (**Fig. 4o-p**).

To determine how a fibroblast-specific modulation could correct myocyte contractile function, single myocyte contraction and Ca^2+^ kinetics were assayed. Unloaded shortening amplitude of intact cardiomyocytes was reduced in I61Q transgenic cardiomyocytes but rescued to WT levels with fibroblast-specific p38 deletion (**Fig. 4q**). This rescue was likely driven by the increased magnitude of the Ca^2+^ transient measured in I61Q-p38^fl/fl^-Tcf21^*iCre*^ cardiomyocytes, which was significantly higher relative to all other experimental genotypes (**Fig. 4r**). To determine if fibroblast-specific p38 deletion also corrects the dilated structural remodeling of the heart, echocardiography was used to measure diastolic chamber dimensions. Here, measurements from I61Q cTnC-p38^fl/fl^-Tcf21^*iCre*^ mice did not show a significant restoration of diastolic chamber dimensions at 4 months of age relative to mice with I61Q cTnC alone, which we ascribe to reduced diastolic tone caused by the I61Q transgene (**Fig. 4s**) (*2*). Since dilated cardiac remodeling is largely a function of serial sarcomere addition which lengthens and thins cardiomyocytes (*42*), morphologic assessment was also performed on cardiomyocytes isolated from the hearts of these experimental mice. I61Q cTnC-p38^fl/fl^-Tcf21^*iCre*^ cardiomyocytes had reduced areas that stemmed from a reduction in cell length when compared to I61Q transgenic cardiomyocytes, which are significantly dilated relative to WT and fibroblast-specific p38 knockout (p38^fl/fl^-Tcf21^*iCre*^) controls (**Fig. 4t-u**). Taken together these data indicate that fibroblast-specific loss of p38 function robustly and simultaneously corrects adaptive remodeling of the ECM and dilated myocyte structure induced by the I61Q mutation in cTnC.

## Conclusion

This study explored the function of fibroblasts as mechanical rheostats within the heart capable of adaptively remodeling the ECM to preserve cardiac function and mechanical homeostasis in response to inherited DCM-linked perturbations in myocyte mechanical function. We believe this is one of a myriad of nested mechanical homeostatic feedback loops guiding organ structure and function in which cells exhibit dynamic reciprocity with their extracellular mechanical environment (*43*–*46*). In DCM, fibroblasts are well-equipped to respond to the contractile insufficiencies of cardiomyocytes, as they are necessarily mechanosensitive to fulfill their role of maintaining tissue integrity (*47*). It is likely that reduced cardiomyocyte tension leads to strain overload as hemodynamic loads on the myocardial wall increase throughout development and disease progression (*48*). Here, both cardiomyocytes and fibroblasts adapted to the pathogenic cTnC variant to preserve the heart’s mechanical integrity and systolic function, where cardiomyocytes altered their morphology and tuned excitation-contraction coupling mechanisms (**Fig. 4q-r, Fig. 4t-u**). Notably, both cardiomyocyte adaptations are highly reversible should the inciting disease stimulus be therapeutically blocked or removed. By contrast cardiac fibroblasts proliferated in response to the I61Q-dependent loss of myocyte tension generation (**Fig. 2f-k**), which is likely a permanent modification given cardiac fibroblasts are resistant to cell death and lack regulatory mechanisms for restricting cell number (*49*–*51*). Hence, the tissue alignment, compaction, and stiffness that resulted from fibroblast proliferation (**Fig. 3a-h**) would likely remain irreversible without a fibroblast-specific therapy that either prevents proliferation or blocks matrix secretion and traction force generation, a result that matches exactly what occurred in this study by silencing p38 activity in cardiac fibroblasts (*41*, *52*). This result is further supported by a recent report that genetic ablation of cardiac fibroblasts during development softens myocardial tissue (*53*). Our finding that the material properties of DCM myocardial tissue is shaped in part by expansion of the cardiac fibroblast population rather than the canonical fibrotic process of fibroblast to myofibroblast transition is critically important to the treatment of non-ischemic DCM, as activated myofibroblast states appear to be transient and unlike changes in fibroblast number these state transitions could resolve or even reverse in response to a DCM specific myocyte targeted therapeutic (*51*). Indeed, first in class therapeutic strategies for DCM like myosin modulators fail to target fibroblast proliferation, which may explain their lukewarm effects on fibrosis (*12*, *54*). It is therefore unlikely that correcting myocyte tension generation alone could reduce fibroblast numbers in the DCM heart unless given at the earliest stage of the disease process. Finally, this study challenges the paradigm that ECM remodeling is secondary to dilated structural remodeling of the myocyte(*3*) and instead supports an active role for fibroblasts in shaping cardiac form and function in DCM, indicating effective therapeutics for this disease will need to address collective cell behaviors rather than singularly restore myocyte function.

## Methods

### Mice

All animal experiments were approved by the University of Washington Institutional Animal Care and Use Committee. I61Q mice were generated as previously described, by mating to a tetracycline transactivator (tTA) line on the FVB/NJ genetic background(*2*). These I61Q tTA mice were further bred onto a line containing LoxP-targeted *Mapk14* (p38^fl/fl^) mice and a tamoxifen regulated Cre recombinase that was knocked into the *Tcf21* locus (Tcf21^iCre^) to generate I61Q cTnC-p38^fl/fl^-Tcf21^*iCre*^ mice (p38 I61Q), which were on mixed genetic background(*41*). Tamoxifen was administered to mice by intraperitoneal injection for 5 consecutive days (400mg/kg body weight in peanut oil), followed by tamoxifen citrate chow ad libitum until a 2 or 4 month experimental endpoint. Echocardiography was performed on a Vevo2100 or Vevo3100 under inhalation isoflurane at heart rates exceeding 350bpm. Invasive hemodynamics on isoflurane-anesthetized mice was performed under heart rates of 420-500bpm using a high-fidelity pressure-volume catheter (1.2F, Transonic) inserted into the left ventricle via the right carotid artery.

### Histology

Fixed cardiac tissues were either processed into paraffin and sectioned (I61Q colony) or cryosectioned in OCT for histologic assessment. Picrosirius red-fast green stained slides were imaged across 6 fields of view at 20x magnification per heart and segmented for collagen content using the color thresholding tool in ImageJ. Whole-heart cross-section images were generated from slide scans obtained by a Hamamtsu Nanozoomer digital pathology system. For fibroblast proliferation and activation, slides were stained with antibodies for αSMA (Sigma A2547, 1:500), phospho-histone H3 (abcam, 1:200), and PDGFRα (1:100 abcam) overnight in staining buffer (1X PBS, 1% BSA, 1% fish skin gelatin), then stained using Alexa Fluor-conjugated secondaries (1:1000 Thermo Fisher) and Hoechst (1:2000 Thermo Fisher) for 90 minutes in staining buffer at room temperature. Stained slides were imaged on a Leica Stellaris 5 confocal microscope under 20x magnification. For quantification, images from six representative regions of interest were obtained at 2x scanner zoom and counted manually blinded to mouse genotype using FIJI(*55*).

### Cardiac Perfusion Decellularization

Freshly harvested hearts were retrograde perfused with a 1% sodium dodecyl sulfate (SDS) solution for 12 hours to decellularize, followed by 1% Triton-X 100 for 1 hour to remove SDS, then rinsed by perfusion with deionized water for 1 hour. Hearts were then transferred to 15mL deionized water, which was refreshed daily for 5 days to ensure complete removal of detergent.

### Multiphoton ECM imaging and structural analysis

Hearts were perfused with 1% agarose and mounted on a 100mm petri dish with the left ventricular free wall facing up, then imaged in whole mount on an Olympus FV1000MP microscope at 25x magnification, using 860nm excitation from a Mai-Tai HP laser (Spectra Physics, 59% power) and a Violet/Green emission filter cube. Z-stacks consisting of 20 images with 1.5-micron step within the LVFW were condensed into maximum intensity projections using ImageJ, then the SHG channel (violet) was quantified for fiber alignment and length using CurveAlign 4.0 beta in CT-FIRE fiber mode (*56*, *57*).

### Cardiac muscle and ECM mechanics

For passive mechanics studies, a Langendorff balloon was inserted into the left ventricle and the heart was perfused with Krebs-Henseleit buffer containing blebbistatin (25μM, Toronto Research Chemicals). The balloon was inflated in 5 microliter steps to 35 μL with 2 minutes of stress relaxation time between each step. This regimen was performed once to precondition the tissue, and then repeated in duplicate for measurements of developed pressure. Following passive muscle measurements, the heart was decellularized as above with the balloon remaining inserted in the left ventricle and the mechanical testing regimen was repeated for the ECM alone. Pressure traces were acquired using LabView and exported to Excel for analysis of developed pressure and curve slope.

### ECM Proteomics

Hearts were perfusion decellularized above, and digested in solution as previously described(*58*). Briefly, samples were first denatured for 2hrs at 37 °C in urea (8M, Fisher) and dithiothreitol (10mM, Thermo Fisher), continuously agitated. Following 30 minutes of alkylation with iodoacetamide (25mM, supplier), samples were then diluted with ammonium bicarbonate (100mM, pH=8.0, Sigma Aldrich), and 2μL PNGase F (500 U/μL, New England Biolabs) was added to deglycosylate the samples over a 2 hour incubation at 37 °C. Samples were then digested by adding 2μL LysC (500ng/μL, Pierce) for 2 hours then 6μL trypsin (100ng/μL New England Biolabs) overnight, both at 37 °C. Trypsin (4μL) was added the next day for 2 hours of additional digestion at 37 °C, then inactivated through addition of 50% trifluoracetic acid (Sigma Aldrich) before samples were clarified through centrifugation (16,000 x g, 5 minutes) and cleaned for liquid chromatography on an MCX column (Waters). For proteomics by data independent acquisition (DIA) mass spectrometry, samples were analyzed at the Nathan Shock Center for Aging proteomics core on a Q Exactive HF Hybrid Quadrupole-Orbitrap Mass Spectrometer (Thermo Fisher) with a Nanoacquity HPLC (Waters). Total ion currents were normalized between samples using the PowerTransformer function of the scikit-learn package in Python, then differential expression between groups was tested by one-way ANOVA(*59*). Figures were generated using Seaborn and matplotlib (*60*).

### Cardiac fibroblast isolation and culture

Primary cardiac fibroblasts were isolated as described previously(*30*). Fibroblasts for RT-PCR and Western blot analyses were negatively sorted on Cd11b microbeads and positively sorted for anti-feeder microbeads through LS columns on a QuadroMACS magnet (Miltenyi Biotec). Fibroblasts for RNASeq, proliferation assays, and engineered tissues were plated on 60mm tissue culture dishes and expanded to the first passage in Dulbecco’s Modified Eagle Medium (DMEM) with 20% fetal bovine serum (FBS) and 1X penicillin/streptomycin solution. Cells in culture were passaged with 0.25% trypsin-EDTA and seeded onto 24-well iBidi μ-plates at a density of 1000 cells/cm^2^ for *in vitro* proliferation studies. The FBS concentration in the media was dropped to 2% upon seeding, and EdU (10μM, Thermo Fisher), dinaciclib (5μM, ApexBio), or ilomastat (10μM, MedChemExpress) were added where indicated. After 24 hours cells were fixed in 4% paraformaldehyde and stained using a Click-It EdU proliferation kit (Thermo Fisher) per manufacturer’s instructions. To screen ECM components, 250,000 WT cardiac fibroblasts were seeded onto an ECM Select Array Kit Ultra-36 (Advanced Biomatrix) and cultured for 24 hours in EdU-containing media as above. Collagen gel compaction was assayed as previously described,(*29*, *61*) with fibroblasts seeded into 1% collagen type I (Advanced Biomatrix) hydrogels at a density of 80,000 cells/mL in a 24-well plate for 24hrs in DMEM with 2% FBS.

### PEG-ECM hydrogels

4-armed PEG_20kDa_-BCN, NHS-Azide, and the MMP-degradable crosslinking peptide N_3_-RGPQGIWGQLPETGGRK(N_3_)-NH_2_ were all synthesized as previously described(*37*, *62*). To generate soluble ECM peptides 4 hearts per genotype were pooled, snap frozen and homogenized by mortar and pestle under liquid nitrogen, lyophilized to a powder, then resuspended at 10mg/mL in a pepsin solution (1mg/mL in 0.1M hydrochloric acid) for 48 hours at room temperature, stirred. Digested ECM was neutralized with the addition of NaOH and re-lyophilized. Digested ECM was resuspended at 25mg/mL in PBS. To azide-functionalize ECM, 2μL of NHS-Azide (60mM in DMSO) was added to 118μL of ECM solution and reacted for 1 hour on ice. Primary cardiac fibroblasts were encapsulated at 10 million cells/mL in gels comprised of 3mM PEG-BCN, 6mM crosslink, 1mM N_3_-GRGDS, and 5mg/mL azide-modified ECM, which were then cultured in DMEM containing 10% FBS and 10μM EdU for 24 hours prior to fixation. Gels were then blocked in PBS containing 0.1M sodium azide to quench any remaining BCN groups in the polymer network and stained as above.

### RNA Sequencing and Analysis

Fibroblasts cultured to 80% confluency were lysed in Trizol (Thermo Fisher) and total RNA was extracted using a Direct-zol RNA Microprep kit, including DNAse treatment (Zymo Research). For RNAseq, RNA integrity was verified using RNA Screentape on a 2200 Tapestation (Agilent) and samples with high RNA integrity (RINe ≥ 7) were submitted to BGI Genomics for RNA sequencing (PE100). Resultant FASTQ files were aligned to the mm10 reference genome using RNA-STAR (*63*), assigned to genes using featurecounts (*64*), and gene transcript counts tested for differential expression using DESeq2(*65*). Differentially expressed genes were tested for pathway enrichment using G:Profiler, and heatmaps were generated in python using the Seaborn package (*60*, *66*). RT-PCR was performed as previously described using the Superscript III First-Strand Synthesis System (Thermo Fisher), iTaq universal SYBR Green Supermix (Bio-Rad) and the primers in **Table S1**(*30*).

### Western Blotting

Magnetically sorted fibroblast pellets were lysed in 120μL of Laemmli Buffer with DTT, of which 30μL was loaded onto a 10% acrylamide gel for sodium dodecyl sulfate polyacrylamide gel electrophoresis (SDS-PAGE) and wet transfer to a polyvinylidene fluoride membrane for immunodetection. Membranes were blocked and immunostained in tris-buffered saline (20mM Tris, 150mM NaCl, pH 7.6) containing 0.1% Tween 20 and 5% nonfat powdered milk. Primary antibodies for phospho-p38 MAPK (Cell Signaling 9211, 1:1000), total p38 MAPK (Cell Signaling 9212, 1:1000), and GAPDH (Fitzgerald 10R-2932, 1:10,000) were incubated overnight at 4°C under gentle agitation. Rabbit or mouse primary antibodies were detected using a horseradish peroxidase-conjugated anti-rabbit IgG (Sigma AP307P, 1:4000) or anti-mouse IgG (Sigma AP308P, 1:4000) secondary antibody for 90 minutes at room temperature, then developed using SuperSignal West Pico PLUS (Thermo Fisher) chemiluminescence substrate.

### Mouse cardiomyocyte isolation and cell culture

For functional measurements, mouse ventricular cardiomyocytes were freshly isolated by Langendorff perfusion with Liberase TM (0.225 mg/mL, Roche) in Krebs-Henseleit buffer (135 mM NaCl, 4.7 mM KCl, 0.6 mM KH_2_PO_4_, 0.6 mM Na_2_HPO_4_, 1.2 mM MgSO_4_, 20 mM HEPES, 10 μM BDM, and 30 mM taurine) as previously described.(*67*) Ventricular cardiomyocytes were mechanically dispersed and filtered through a 200 μm nylon mesh then allowed to sediment for 5-10 minutes. Sedimentation was repeated three times using increasing [Ca^2+^] from 0.125 to 0.25 to 0.5 mmol/L. Cardiomyocytes were plated on laminin-coated coverslips in Tyrode’s solution (137 mM NaCl, 5.4 mM KCl, 0.5 mM MgCl_2_, 1.2 mM CaCl_2_•2H_2_O, 10 mM HEPES, and 5 mM Glucose, pH 7.4) for 1 hour at 37 °C prior to functional measurements. For myocyte morphology measurements, cardiomyocytes were similarly isolated and plated with buffers containing 25 μM blebbistatin and subsequently fixed with 4% PFA at room temperature for 15 minutes.

### Measurements of cardiomyocyte contractility and calcium transients

Sarcomere measurements were obtained from isolated cardiomyocytes using the IonOptix™ SarcLen Sarcomere Length Acquisition Module with a MyoCam-S3 digital camera (Ionoptix Co.) attached to an Olympus uWD 40 inverted microscope. For these measurements cardiomyocytes were bathed in 1.2 mM Ca^2+^ Tyrode’s solution and kept at 37 °C. To jumpstart pacing, cardiomyocytes were stimulated with frequencies varying from 0.5, 1.0, and 1.5 Hz at 10 V for a minimum of 10 contractions at each frequency. Sarcomere lengths were then measured in real time at a frequency of 1 Hz and averaged across 10-15 contraction cycles. Separate coverslips were treated with 1 μM Fura-2–acetoxymethyl ester to measure calcium transients. Blinded analysis was performed using the IonWizard software. Statistical analyses were performed on individual myocyte measurements (n ~ 20 cardiomyocytes/mouse; n=3-4). Significance was determined using Student’s t-test. For myocyte geometry quantification approximately 50 cells per mouse were manually traced using FIJI.

### Rat cardiomyocyte isolation and EHT experiments

Freshly isolated neonatal rat cardiomyocytes and fibroblasts were seeded into 100 μL fibrin EHTs containing 1 million cells per tissue between a pair of flexible and rigid PDMS posts that were 12 mm in length and 1.5 mm in diameter within a 24-well plate, as previously described (*68*). EHTs were polymerized for 85 minutes, then demolded and immersed in plating media [4:1 DMEM:Medium 199 (M199), 10% horse serum, 5% FBS, 100 U/mL penicillin streptomycin (pen-strep)] containing AdGFP or AdI61Q at a multiplicity of infection of 200. After 24 hours, EHTs were switched to maintenance medium consisting of 1:1 DMEM:M199 containing 5% FBS, 100 U/mL pen-strep, 5 g/L 6-aminocaproic acid, 1X insulin-transferrin-selenium, and 0.1% chemically defined lipid concentrate, which was thenceforth swapped every other day until the 14-day experimental endpoint. EHTs were then bathed in Tyrode’s buffer equilibrated to 37 °C for contractile analysis as previously described (*68*). Briefly, EHTs were paced at 1 Hz by a custom 24-well plate pacing apparatus with carbon electrodes biphasically stimulated (5V/cm, 10ms duration) with a medical stimulator (Astro Med Grass Stimulator, Model S88X) while imaged. Brightfield videos of PDMS post deflection during EHT contraction were taken at 66.67 frames per second on a Nikon TEi epi-fluorescent microscope under 2x magnification. Deflection of the flexible post relative to the rigid post was tracked using a custom MATLAB script in order to calculate twitch force, and tension index as the area under the twitch curve. Following functional measurements, EHTs were fixed in ice-cold 4% PFA for 1 hour and stained with anti-FLAG (Sigma, 1:1000), Alexa Fluor 568-conjugated wheat germ agglutinin (Thermo Fisher 1:1000), and Hoechst 33342 (Thermo Fisher, 1:1000). For alignment, 8 ROIs per wheat-germ stained EHT were confocally imaged in whole mount at 20x magnification on a Leica Stellaris 5 confocal microscope and analyzed using the Directionality plugin in FIJI. Alignment coefficient was calculated as the amount divided by the dispersion of directionality.

### Intact and skinned muscle mechanics

Hearts were quickly removed via thoracotomy and rinsed in oxygenated modified Krebs buffer containing 118.5 mM NaCl, 5 mM KCl, 1.2 mM MgSO_4_, 2 mM NaH_2_PO_4_, 25 mM NaHCO_3_, 1.8 mM CaCl_2_, and 10 mM glucose. Hearts were then perfused and dissected in oxygenated modified Krebs with 0.1 mM CaCl_2_ and 20 mM 2,3-butanedione 2-monoxime (BDM) to limit contraction and damage during tissue dissection.

For demembranated tissue mechanics, dissected hearts were permeabilized in a glycerol relaxing solution containing 100 mM KCl, 10 mM MOPS, 5 mM K_2_EGTA, 9 mM MgCl_2_ and 5 mM Na_2_ATP (pH 7.0), 1% (by vol) Triton X-100, 1% protease inhibitor (Sigma P8340), and 50% (by vol) glycerol at 4°C overnight then stored in fresh solution without Triton X-100 for storage at −20°C. Briefly, right ventricular trabeculae were dissected and mounted between a force transducer and motor, and sarcomere length (SL) was set to ~2.3 μm, as previously described.(*5*) Experiments were conducted in a physiological solution (15°C, pH 7.0) containing a range of pCa (= –log[Ca^2+^]) from 9.0 to 4.0. Force and k_tr_ (rate of tension redevelopment) were collected at each pCa and analyzed with custom using LabView software.

For intact twitch measurements unbranched, intact trabeculae were dissected from the right ventricular wall and mounted between a force transducer (Cambridge Technology, Inc., model 400A) and a rigid post, as previously described. The tissue was then submerged in a custom experimental chamber that was continuously perfused with oxygenated modified Krebs buffer (1.8 mM CaCl_2_) at 33°C. After a ~20min equilibration and washout at 0.5 Hz pacing, optimal length was set to ~2.3 μm SL and tissue was paced at 1 Hz. 30 second traces were recorded on custom LabView software and were analyzed with custom code written using MATLAB software (Mathworks).

## Acknowledgements

The authors would like to acknowledge the support of Mike MacCoss and the proteomics core at the University of Washington (UW) Nathan Shock Center for Aging, Dale Hailey and the Garvey Imaging core at the UW Institute for Stem Cell and Regenerative Medicine (ISCRM), and Brian Johnson and the UW Histology and Imaging Core. Thanks to Rong Tian for use of critical instrumentation, Saffie Mohran for insightful discussion, Emily Olszewski and other members of the Davis Lab for their insights and experimental assistance.

## Author Contributions

RCB, IMR, KAZ, AG, LB, DB, TM, KK, GF, AM, JG, FK, EP, WAL, and JD performed experiments and analyzed results. Experiments were conceived by RCB, IMR, KAZ, TM, KK, WL, MR, FMH, NJS, CAD, and JD, RCB, MR, NJS, CAD, and JD wrote and reviewed the manuscript.

## Supplementary Materials

Materials and Methods

Figs S1 to S7

Table S1

Data S1

## References

1. R. Yotti, C. E. Seidman, J. G. Seidman, Advances in the Genetic Basis and Pathogenesis of Sarcomere Cardiomyopathies. Annu. Rev. Genomics Hum. Genet. 20, 129–153 (2019).

2. J. Davis, L. C. Davis, R. N. Correll, C. A. Makarewich, J. A. Schwanekamp, F. Moussavi-Harami, D. Wang, A. J. York, H. Wu, S. R. Houser, C. E. Seidman, J. G. Seidman, M. Regnier, J. M. Metzger, J. C. Wu, J. D. Molkentin, A Tension-Based Model Distinguishes Hypertrophic versus Dilated Cardiomyopathy. Cell. 165, 1147–1159 (2016).

3. T. R. Eijgenraam, H. H. W. Silljé, R. A. de Boer, Current understanding of fibrosis in genetic cardiomyopathies. Trends Cardiovasc. Med. 30, 353–361 (2020).

4. R. C. Bretherton, D. Bugg, E. O. Olszewski, J. Davis, Regulators of Cardiac Fibroblast Cell State. Matrix Biol. 91–92, 117–135 (2020).

5. J. D. Powers, K. B. Kooiker, A. B. Mason, A. E. Teitgen, G. V. Flint, J. C. Tardiff, S. D. Schwartz, A. D. McCulloch, M. Regnier, J. Davis, F. Moussavi-Harami, Modulating the tension-time integral of the cardiac twitch prevents dilated cardiomyopathy in murine hearts. JCI Insight. 5(2020), doi:10.1172/jci.insight.142446.

6. R. E. Hershberger, D. J. Hedges, A. Morales, Dilated cardiomyopathy: the complexity of a diverse genetic architecture. Nat. Rev. Cardiol. 10, 531–547 (2013).

7. E. M. Mcnally, J. R. Golbus, M. J. Puckelwartz, Genetic mutations and mechanisms in dilated cardiomyopathy. J Clin Invest. 123, 19–26 (2013).

8. G. W. Dec, V. Fuster, Idiopathic Dilated Cardiomyopathy. N. Engl. J. Med. 331, 1564–1575 (1994).

9. B. P. Halliday, A. J. Baksi, A. Gulati, A. Ali, S. Newsome, C. Izgi, M. Arzanauskaite, A. Lota, U. Tayal, V. S. Vassiliou, J. Gregson, F. Alpendurada, M. P. Frenneaux, S. A. Cook, J. G. F. Cleland, D. J. Pennell, S. K. Prasad, Outcome in Dilated Cardiomyopathy Related to the Extent, Location, and Pattern of Late Gadolinium Enhancement. JACC Cardiovasc. Imaging. 12, 1645–1655 (2019).

10. A. Gulati, A. G. Japp, S. Raza, B. P. Halliday, D. A. Jones, S. Newsome, N. A. Ismail, K. Morarji, J. Khwaja, N. Spath, C. Shakespeare, P. R. Kalra, G. Lloyd, A. Mathur, J. G. F. Cleland, M. R. Cowie, R. G. Assomull, D. J. Pennell, T. F. Ismail, S. K. Prasad, Absence of Myocardial Fibrosis Predicts Favorable Long-Term Survival in New-Onset Heart Failure. Circ. Cardiovasc. imaging. 11, e007722 (2018).

11. A. A. Mandawat, P. Chattranukulchai, A. A. Mandawat, A. J. Blood, S. Ambati, B. Hayes, W. Rehwald, H. W. Kim, J. F. Heitner, D. J. Shah, I. Klem, Progression of Myocardial Fibrosis in Nonischemic DCM and Association With Mortality and Heart Failure Outcomes. JACC Cardiovasc. Imaging. 14, 1338–1350 (2021).

12. J. R. Teerlink, R. Diaz, G. M. Felker, J. J. V. McMurray, M. Metra, S. D. Solomon, K. F. Adams, I. Anand, A. Arias-Mendoza, T. Biering-Sørensen, M. Böhm, D. Bonderman, J. G. F. Cleland, R. Corbalan, M. G. Crespo-Leiro, U. Dahlström, L. E. Echeverria, J. C. Fang, G. Filippatos, C. Fonseca, E. Goncalvesova, A. R. Goudev, J. G. Howlett, D. E. Lanfear, J. Li, M. Lund, P. Macdonald, V. Mareev, S. Momomura, E. O’Meara, A. Parkhomenko, P. Ponikowski, F. J. A. Ramires, P. Serpytis, K. Sliwa, J. Spinar, T. M. Suter, J. Tomcsanyi, H. Vandekerckhove, D. Vinereanu, A. A. Voors, M. B. Yilmaz, F. Zannad, L. Sharpsten, J. C. Legg, C. Varin, N. Honarpour, S. A. Abbasi, F. I. Malik, C. E. Kurtz, Cardiac Myosin Activation with Omecamtiv Mecarbil in Systolic Heart Failure. N. Engl. J. Med. 384, 105–116 (2021).

13. O. Kanisicak, H. Khalil, M. J. Ivey, J. Karch, B. D. Maliken, R. N. Correll, M. J. Brody, S.-C. J Lin, B. J. Aronow, M. D. Tallquist, J. D. Molkentin, Genetic lineage tracing defines myofibroblast origin and function in the injured heart. Nat. Commun. 7, 12260 (2016).

14. A. Acharya, S. T. Baek, G. Huang, B. Eskiocak, S. Goetsch, C. Y. Sung, S. Banfi, M. F. Sauer, G. S. Olsen, J. S. Duffield, E. N. Olson, M. D. Tallquist, The bHLH transcription factor Tcf21 is required for lineage-specific EMT of cardiac fibroblast progenitors. Development. 139, 2139–2149 (2012).

15. T. Moore-morris, N. Guimarães-camboa, I. Banerjee, A. C. Zambon, T. Kisseleva, A. Velayoudon, W. B. Stallcup, Y. Gu, N. D. Dalton, M. Cedenilla, R. Gomez-amaro, B. Zhou, D. A. Brenner, K. L. Peterson, J. Chen, S. M. Evans, Resident fibroblast lineages mediate pressure overload-induced cardiac fibrosis. J. Clin. Invest. 124, 1–14 (2014).

16. S. R. Ali, S. Ranjbarvaziri, M. Talkhabi, P. Zhao, A. Subat, A. Hojjat, P. Kamran, A. M. S. Müller, K. S. Volz, Z. Tang, K. Red-Horse, R. Ardehali, Developmental heterogeneity of cardiac fibroblasts does not predict pathological proliferation and activation. Circ. Res. 115 625–635 (2014).

17. J. J. Saucerman, P. M. Tan, K. S. Buchholz, A. D. McCulloch, J. H. Omens, Mechanical regulation of gene expression in cardiac myocytes and fibroblasts. Nat. Rev. Cardiol. 2019 166. 16, 361–378 (2019).

18. K. L. Kreutziger, N. Piroddi, J. T. McMichael, C. Tesi, C. Poggesi, M. Regnier, Calcium binding kinetics of troponin C strongly modulate cooperative activation and tension kinetics in cardiac muscle. J. Mol. Cell. Cardiol. 50, 165–174 (2011).

19. S. B. Tikunova, J. P. Davis, Designing calcium-sensitizing mutations in the regulatory domain of cardiac troponin C. J. Biol. Chem. 279, 35341–35352 (2004).

20. L. J. Dooling, K. Saini, A. A. Anlaş, D. E. Discher, Tissue mechanics coevolves with fibrillar matrisomes in healthy and fibrotic tissues. Matrix Biol. 111, 153–188 (2022).

21. R. O. Hynes, A. Naba, Overview of the matrisome--an inventory of extracellular matrix constituents and functions. Cold Spring Harb. Perspect. Biol. 4, a004903 (2012).

22. M. Cescon, F. Gattazzo, P. Chen, P. Bonaldo, Collagen VI at a glance. J. Cell Sci. 128, 3525–3531 (2015).

23. J. A. Chirinos, L. Zhao, A. L. Reese-Petersen, J. B. Cohen, F. Genovese, A. M. Richards, R. N. Doughty, J. Díez, A. González, R. Querejeta, P. Zamani, J. Nuñez, Z. Wang, C. Ebert, K. Kammerhoff, J. Maranville, M. Basso, C. Qian, D. G. K. Rasmussen, P. H. Schafer, D. Seiffert, M. A. Karsdal, D. A. Gordon, F. Ramirez-Valle, T. P. Cappola, Endotrophin, a Collagen VI Formation–Derived Peptide, in Heart Failure. NEJM Evid. 1(2022).

24. P. Fratzl, Collagen: Structure and mechanics, an introduction. Collagen Struct. Mech., 1–13 (2008).

25. G. M. Fomovsky, S. Thomopoulos, J. W. Holmes, Contribution of extracellular matrix to the mechanical properties of the heart. J. Mol. Cell. Cardiol. 48, 490–496 (2010).

26. N. Hamdani, M. Herwig, W. A. Linke, Tampering with springs: phosphorylation of titin affecting the mechanical function of cardiomyocytes. Biophys. Rev. 9, 225 (2017).

27. M. M. Lewinter, H. L. Granzier, Cardiac Titin and Heart Disease. J. Cardiovasc. Pharmacol. 63, 207 (2014).

28. J. Davis, J. D. Molkentin, Myofibroblasts: Trust your heart and let fate decide. J. Mol. Cell. Cardiol. 70(2014).

29. J. Davis, N. Salomonis, N. Ghearing, S.-C. J. Lin, J. Q. Kwong, A. Mohan, M. S. Swanson, J. D. Molkentin, MBNL1-mediated regulation of differentiation RNAs promotes myofibroblast transformation and the fibrotic response. Nat. Commun. 6, 10084 (2015).

30. D. Bugg, L. R. J. Bailey, R. C. Bretherton, K. E. Beach, I. M. Reichardt, K. Z. Robeson, C. Reese, J. Gunaje, G. Flint, C. A. DeForest, A. Stempien-Otero, J. Davis, MBNL1 drives dynamic transitions between fibroblasts and myofibroblasts in cardiac wound healing. Cell Stem Cell. 0(2022), doi:10.1016/J.STEM.2022.01.012.

31. D. A. Skelly, G. T. Squiers, M. A. McLellan, M. T. Bolisetty, P. Robson, N. A. Rosenthal, R. Pinto, Single-Cell Transcriptional Profiling Reveals Cellular Diversity and Intercommunication in the Mouse Heart. Cell Rep. 22, 600–610 (2018).

32. N. Farbehi, R. Patrick, A. Dorison, M. Xaymardan, V. Janbandhu, K. Wystub-Lis, J. Wk Ho, R. E. Nordon, R. P. Harvey, Single-cell expression profiling reveals dynamic flux of cardiac stromal, vascular and immune cells in health and injury (2019).

33. M. B. Buechler, R. N. Pradhan, A. T. Krishnamurty, C. Cox, A. K. Calviello, A. W. Wang, Y. A. Yang, L. Tam, R. Caothien, M. Roose-Girma, Z. Modrusan, J. R. Arron, R. Bourgon, S. Müller, S. J. Turley, Cross-tissue organization of the fibroblast lineage. Nat. 593, 575–579 (2021).

34. U. Raudvere, L. Kolberg, I. Kuzmin, T. Arak, P. Adler, H. Peterson, J. Vilo, G:Profiler: A web server for functional enrichment analysis and conversions of gene lists (2019 update). Nucleic Acids Res. 47, W191–W198 (2019).

35. C. Williams, K. P. Quinn, I. Georgakoudi, L. D. Black, Young developmental age cardiac extracellular matrix promotes the expansion of neonatal cardiomyocytes in vitro. Acta Biomater. 10, 194–204 (2014).

36. C. A. DeForest, B. D. Polizzotti, K. S. Anseth, Sequential click reactions for synthesizing and patterning three-dimensional cell microenvironments. Nat. Mater. 8, 659–664 (2009).

37. C. A. DeForest, D. A. Tirrell, A photoreversible protein-patterning approach for guiding stem cell fate in three-dimensional gels. Nat. Mater. 14, 523–531 (2015).

38. S. Checa, M. K. Rausch, A. Petersen, E. Kuhl, G. N. Duda, The emergence of extracellular matrix mechanics and cell traction forces as important regulators of cellular self-organization. Biomech. Model. Mechanobiol. 2014 141. 14, 1–13 (2014).

39. D. Bugg, R. Bretherton, P. Kim, E. Olszewski, A. Nagle, A. E. Schumacher, N. Chu, J. Gunaje, C. A. DeForest, K. Stevens, D.-H. Kim, J. Davis, Infarct Collagen Topography Regulates Fibroblast Fate via p38-Yes-Associated Protein Transcriptional Enhanced Associate Domain Signals. Circ. Res. 127(2020).

40. Turner, Blythe, Cardiac Fibroblast p38 MAPK: A Critical Regulator of Myocardial Remodeling. J. Cardiovasc. Dev. Dis. 6, 27 (2019).

41. J. D. Molkentin, D. Bugg, N. Ghearing, L. E. Dorn, P. Kim, M. A. Sargent, J. Gunaje, K. Otsu, J. Davis, Fibroblast-Specific Genetic Manipulation of p38 MAPK in vivo Reveals its Central Regulatory Role in Fibrosis. Circulation. 136, 549–561 (2017).

42. I. Kehat, J. Davis, M. Tiburcy, F. Accornero, M. K. Saba-El-Leil, M. Maillet, A. J. York, J. N. Lorenz, W. H. Zimmermann, S. Meloche, J. D. Molkentin, Extracellular Signal-Regulated Kinases 1 and 2 Regulate the Balance Between Eccentric and Concentric Cardiac Growth. Circ. Res. 108, 176–183 (2011).

43. C. J. Chan, C. P. Heisenberg, T. Hiiragi, Coordination of Morphogenesis and Cell-Fate Specification in Development. Curr. Biol. 27, R1024–R1035 (2017).

44. C. P. Heisenberg, Y. Bellaïche, Forces in tissue morphogenesis and patterning. Cell. 153, 948 (2013).

45. D. Gilmour, M. Rembold, M. Leptin, From morphogen to morphogenesis and back. Nature. 541, 311–320 (2017).

46. N. I. Petridou, Z. Spiró, C. P. Heisenberg, Multiscale force sensing in development. Nat. Cell Biol. 19, 581–588 (2017).

47. M. Pesce, G. N. Duda, G. Forte, H. Girao, A. Raya, P. Roca-Cusachs, J. P. G. Sluijter, C. Tschöpe, S. Van Linthout, Cardiac fibroblasts and mechanosensation in heart development, health and disease. Nat. Rev. Cardiol. 2022, 1–16 (2022).

48. A. G. Rodriguez, S. J. Han, M. Regnier, N. J. Sniadecki, Substrate Stiffness Increases Twitch Power of Neonatal Cardiomyocytes in Correlation with Changes in Myofibril Structure and Intracellular Calcium. Biophys. J. 101, 2455–2464 (2011).

49. J. T. Kuwabara, A. Hara, S. Bhutada, G. S. Gojanovich, J. Chen, K. Hokutan, V. Shettigar, A. Y. Lee, L. P. Deangelo, J. R. Heckl, J. R. Jahansooz, D. K. Tacdol, M. T. Ziolo, S. S. Apte, M. D. Tallquist, Consequences of PDGFRα+ fibroblast reduction in adult murine hearts. Elife. 11(2022), doi:10.7554/ELIFE.69854.

50. M. J. Ivey, J. T. Kuwabara, K. L. Riggsbee, M. D. Tallquist, Platelet-derived growth factor receptor-α is essential for cardiac fibroblast survival. Am. J. Physiol. Circ. Physiol. 317, H330–H344 (2019).

51. X. Fu, H. Khalil, O. Kanisicak, J. G. Boyer, R. J. Vagnozzi, B. D. Maliken, M. A. Sargent, V. Prasad, I. Valiente-Alandi, B. C. Blaxall, J. D. Molkentin, Specialized fibroblast differentiated states underlie scar formation in the infarcted mouse heart. J. Clin. Invest. 128, 2127–2143 (2018).

52. D. Bugg, R. C. Bretherton, P. Kim, E. Olszewski, A. Nagle, A. E. A. E. Schumacher, N. Chu, J. Gunaje, C. A. C. A. DeForest, K. Stevens, D.-H. D.-H. Kim, J. M. Davis, Infarct Collagen Topography Regulates Fibroblast Fate Via p38-Yap-TEAD Signals. Circ. Res. 127(2020).

53. J. T. Kuwabara, A. Hara, J. R. Heckl, B. Peña, S. Bhutada, R. DeMaris, M. J. Ivey, L. P. DeAngelo, X. Liu, J. Park, J. R. Jahansooz, L. Mestroni, T. A. McKinsey, S. S. Apte, M. D. Tallquist, Regulation of extracellular matrix composition by fibroblasts during perinatal cardiac maturation. J. Mol. Cell. Cardiol. (2022).

54. S. Saberi, N. Cardim, M. Yamani, J. Schulz-Menger, W. Li, V. Florea, A. J. Sehnert, R. Y. Kwong, M. Jerosch-Herold, A. Masri, A. Owens, N. K. Lakdawala, C. M. Kramer, M. Sherrid, T. Seidler, A. Wang, F. Sedaghat-Hamedani, B. Meder, O. Havakuk, D. Jacoby, Mavacamten Favorably Impacts Cardiac Structure in Obstructive Hypertrophic Cardiomyopathy: EXPLORER-HCM Cardiac Magnetic Resonance Substudy Analysis. Circulation. 143, 606–608 (2021).

55. J. Schindelin, I. Arganda-Carreras, E. Frise, V. Kaynig, M. Longair, T. Pietzsch, S. Preibisch, C. Rueden, S. Saalfeld, B. Schmid, J.-Y. Tinevez, D. J. White, V. Hartenstein, K. Eliceiri, P. Tomancak, A. Cardona, Fiji: an open-source platform for biological-image analysis. Nat. Methods. 9, 676–682 (2012).

56. J. S. Bredfeldt, Y. Liu, C. A. Pehlke, M. W. Conklin, J. M. Szulczewski, D. R. Inman, P. J. Keely, R. D. Nowak, T. R. Mackie, K. W. Eliceiri, Computational segmentation of collagen fibers from second-harmonic generation images of breast cancer. J. Biomed. Opt. 19, 016007 (2014).

57. Y. Liu, A. Keikhosravi, G. S. Mehta, C. R. Drifka, K. W. Eliceiri, Methods for quantifying fibrillar collagen alignment. Methods Mol. Biol. 1627, 429–451 (2017).

58. A. Naba, K. R. Clauser, R. O. Hynes, Enrichment of Extracellular Matrix Proteins from Tissues and Digestion into Peptides for Mass Spectrometry Analysis. JoVE (Journal Vis. Exp. 2015, e53057 (2015).

59. F. Pedregosa, V. Michel, O. Grisel, M. Blondel, P. Prettenhofer, R. Weiss, J. Vanderplas, D. Cournapeau, F. Pedregosa, G. Varoquaux, A. Gramfort, B. Thirion, O. Grisel, V. Dubourg, A. Passos, M. Brucher, M. Perrot, E. Duschenay, “Scikit-learn: Machine Learning in Python” (2011), (available at http://scikit-learn.sourceforge.net.).

60. M. Waskom, Seaborn (2020),, doi:10.5281/zenodo.592845.

61. P. Ngo, P. Ramalingam, J. A. Phillips, G. T. Furuta, “Collagen Gel Contraction Assay” in Cell-Cell Interactions in Health and Disease (Humana Press, New Jersey, 2006; http://link.springer.com/10.1385/1-59745-113-4:103), pp. 103–110.

62. U. N. Lee, J. H. Day, A. J. Haack, R. C. Bretherton, W. Lu, C. A. DeForest, A. B. Theberge, E. Berthier, Layer-by-layer fabrication of 3D hydrogel structures using open microfluidics. Lab Chip. 20, 525–536 (2020).

63. A. Dobin, C. A. Davis, F. Schlesinger, J. Drenkow, C. Zaleski, S. Jha, P. Batut, M. Chaisson, T. R. Gingeras, STAR: Ultrafast universal RNA-seq aligner. Bioinformatics. 29, 15–21 (2013).

64. Y. Liao, G. K. Smyth, W. Shi, Sequence analysis featureCounts: an efficient general purpose program for assigning sequence reads to genomic features. 30, 923–930 (2014).

65. M. I. Love, W. Huber, S. Anders, Moderated estimation of fold change and dispersion for RNA-seq data with DESeq2. Genome Biol. 15, 550 (2014).

66. U. Raudvere, L. Kolberg, I. Kuzmin, T. Arak, P. Adler, H. Peterson, J. Vilo, g:Profiler: a web server for functional enrichment analysis and conversions of gene lists (2019 update). Nucleic Acids Res. 47, W191–W198 (2019).

67. B. Hegyi, J. M. Borst, L. R. J. Bailey, E. Y. Shen, A. J. Lucena, M. F. Navedo, J. Bossuyt, D. M. Bers, Hyperglycemia regulates cardiac K+ channels via O-GlcNAc-CaMKII and NOX2-ROS-PKC pathways. Basic Res. Cardiol. 115, 1–19 (2020).

68. S. Bremner, A. J. Goldstein, T. Higashi, N. J. Sniadecki, Engineered Heart Tissues for Contractile, Structural, and Transcriptional Assessment of Human Pluripotent Stem Cell-Derived Cardiomyocytes in a Three-Dimensional, Auxotonic Environment. Methods Mol. Biol. 2485, 87–97 (2022).

