## Supplemental Material for "Correcting dilated cardiomyopathy with fibroblast-targeted p38 deficiency"

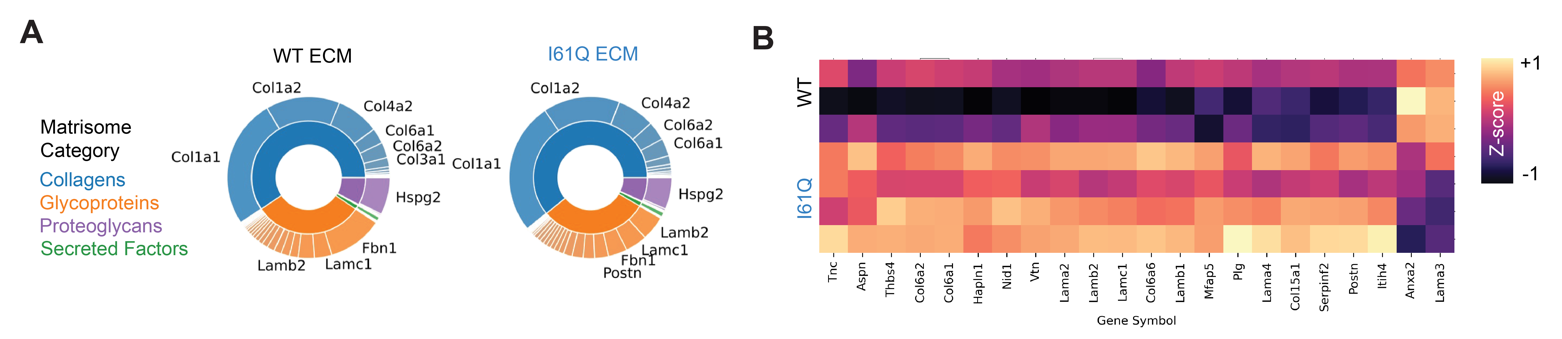

**Fig. S1. Matrisomal analysis of the I61Q matrix in 4-month-old mice.** **(A)** Pie charts showing the relative abundance of matrisomal proteins colored by category, with the top 10 most abundant proteins labeled. **(B)** Heatmap of significantly differentially regulated matrisome proteins color coded by z-score.

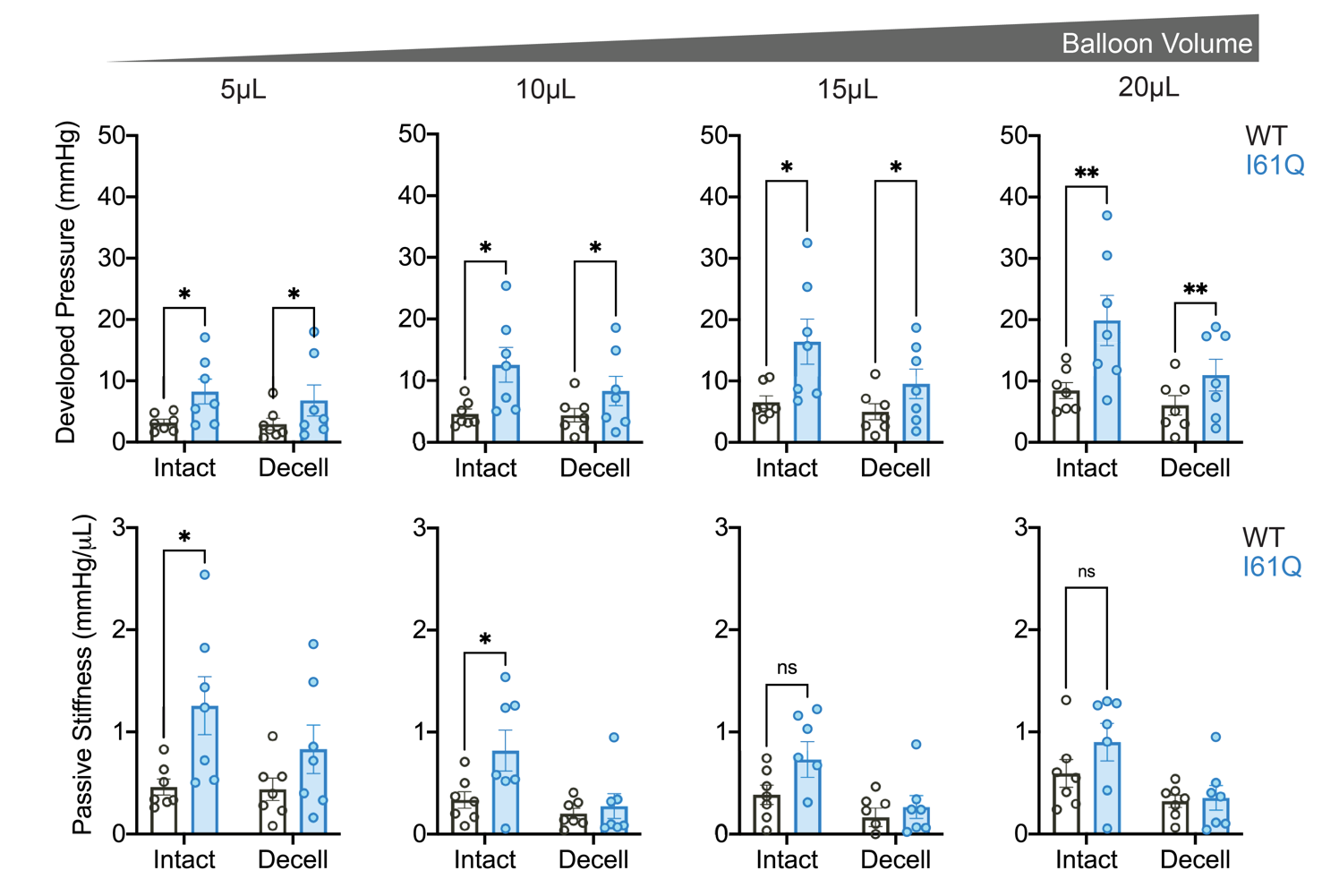

**Fig. S2. Mechanical properties of myocardium and matrix measured by pressure-volume balloon.** Developed balloon pressure (top) and passive stiffness measured as the slope of the pressure volume curve varying balloon volume (bottom). ns = nonsignificant, *p<0.05,**p<0.01 by 2-way ANOVA with Holm-Sidak’s multiple comparisons test.

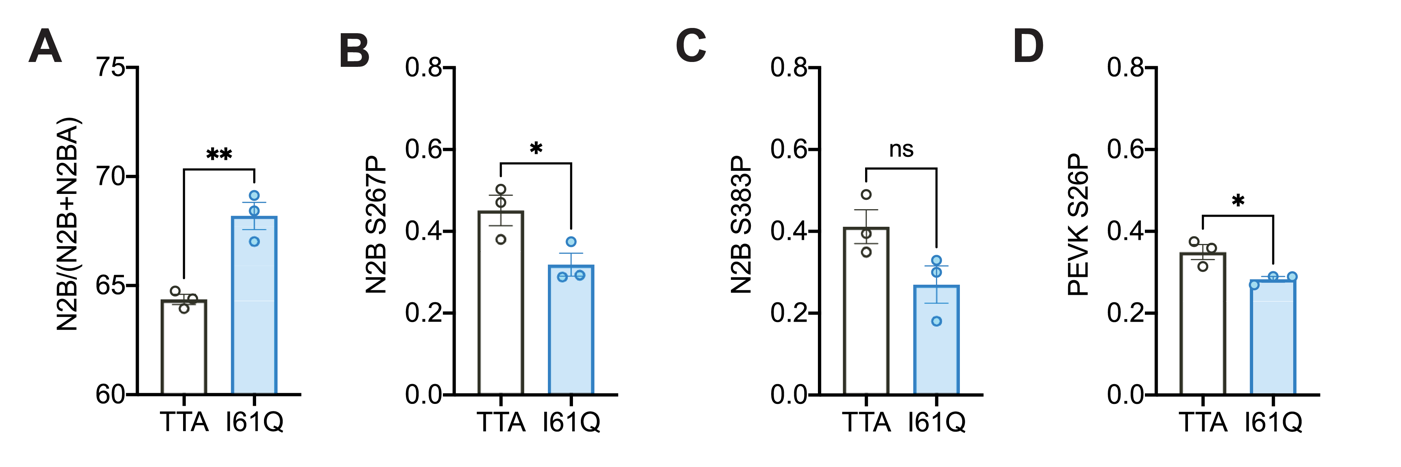

**Fig. S3. Titin isoforms and phosphorylation status shift in 2-month-old I61Q mice** **(A)** Isoform analysis reveals a shift towards the noncompliant N2B titin isoform in I61Q myocardium. **(B)** Reduced phosphorylation of N2B serine 267 was observed in I61Q hearts by western blot. **(C)** phosphorylation of N2B serine 383 was observed. **(D)** phosphorylation of serine 267 in the PEVK region. ns = nonsignificant, *p<0.05,**p<0.01 by unpaired t-test.

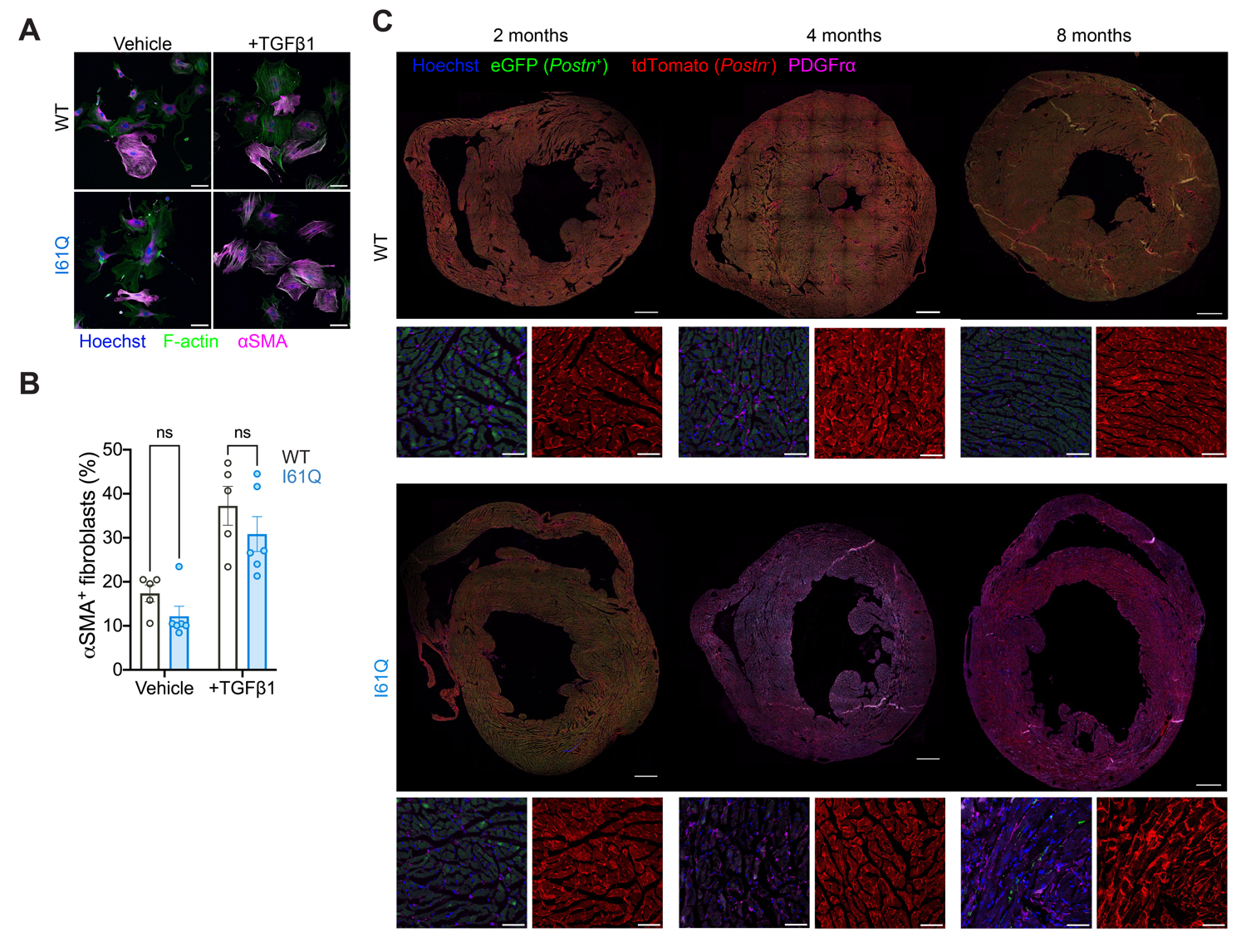

**Fig. S4. Fibroblast activation does not underly cardiac fibrosis in I61Q hearts.** **(A)** Representative images (scale 50μm) and **(B)** quantification of fibroblast-to-myocyte conversion by aSMA immunocytochemistry. **(C)** Representative cross sections (scale 1mm) and 20x regions of interest (scale 50μm) from Postn^mT/mG^ I61Q TTA hearts and WT controls. ns = nonsignificant by 2-way ANOVA with Holm-Sidak’s multiple comparisons test.

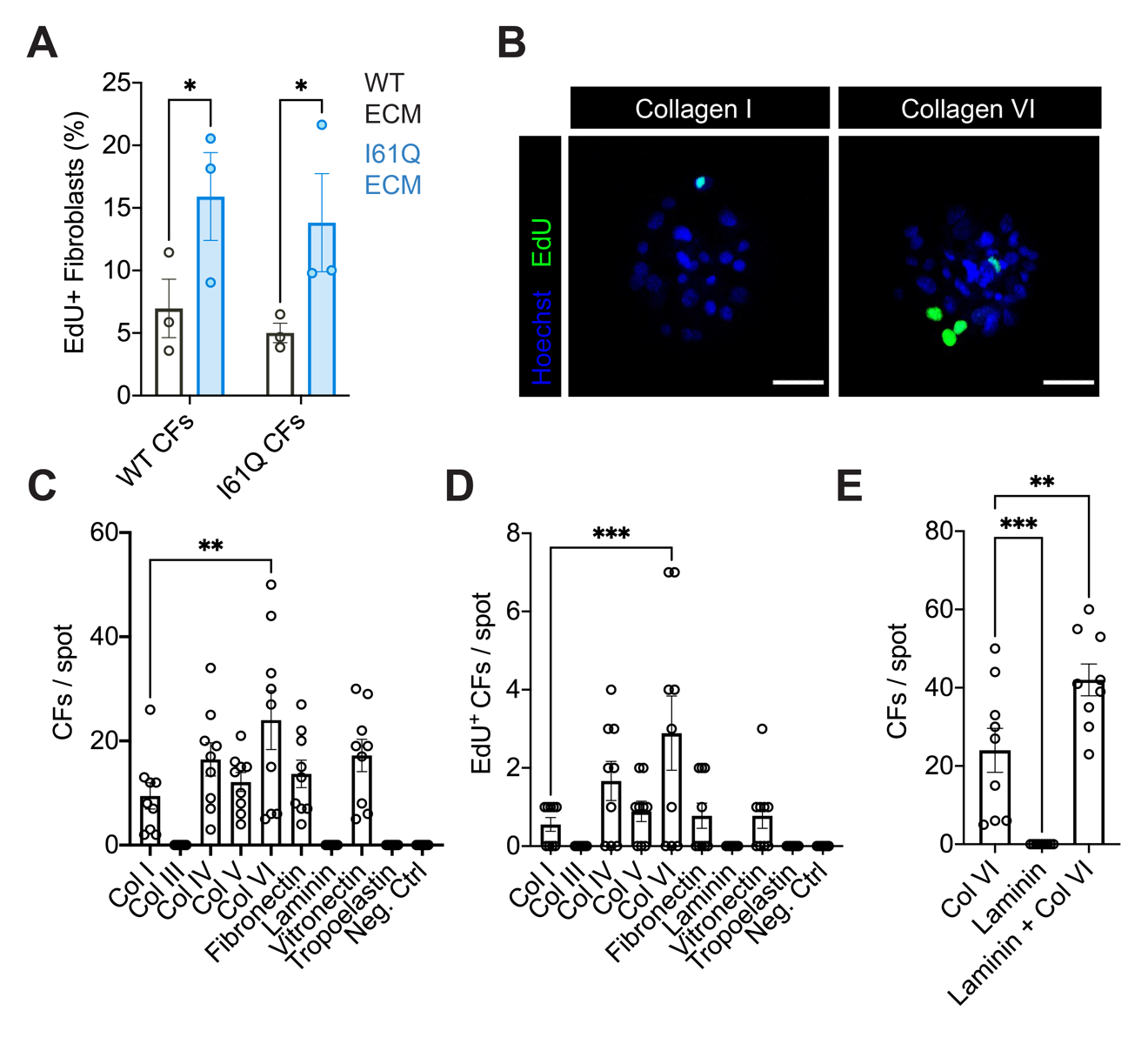

**Fig. S5. ECM cues driving cardiac fibroblast adhesion and proliferation. (A)** Proliferation by EdU assay of WT and I61Q fibroblasts seeded within PEG gels decorated with ECM peptides from WT and I61Q hearts. **(B)** Representative images of cardiac fibroblasts seeded onto the ECM screening array stained for EdU to mark proliferating cells. **(C)** Fibroblast counts and **(D)** EdU+ fibroblast counts on ECM-coated microspots. **(E)** Fibroblast counts on spots containing collagen VI only, laminin only, or both laminin and collagen VI. *p<0.05,**p<0.01,***p<0.005 by either 2-way ANOVA (**A)** or one-way ANOVA (**C-E**) with Holm-Sidak’s multiple comparisons test.

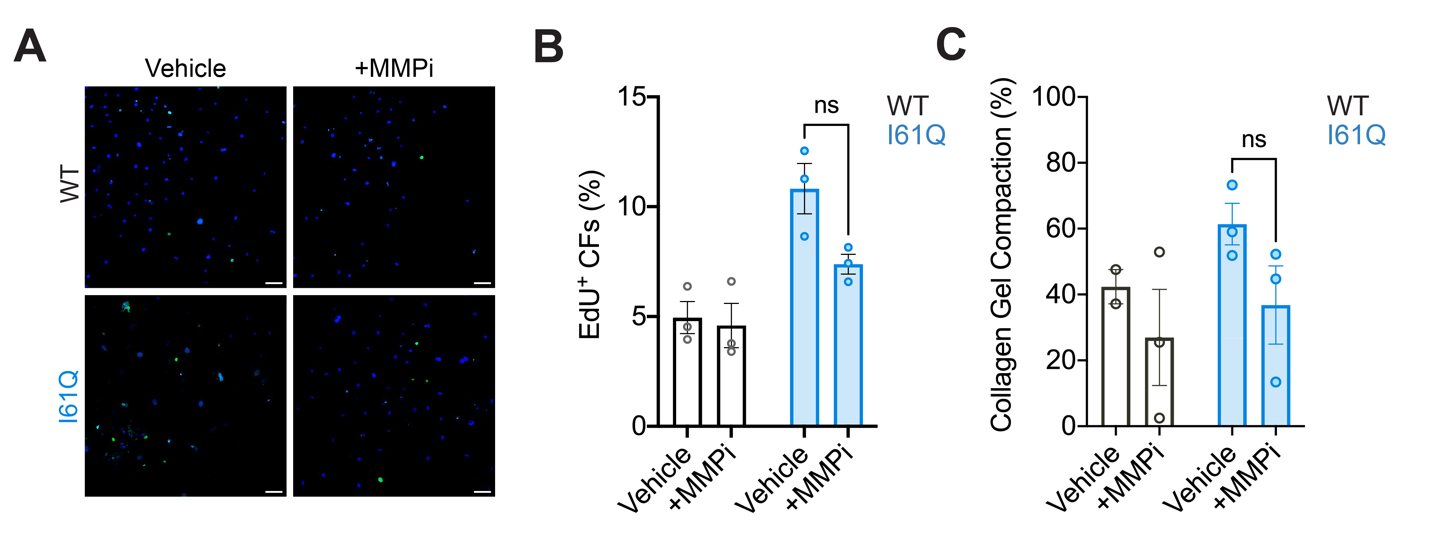

**Fig. S6. MMP inhibition does not significantly reduce cardiac fibroblast proliferation or collagen gel compaction. (A)** Representative EdU-stained immunofluorescent images (scale 50μm) and **(B)** quantification of WT and I61Q fibroblasts treated with vehicle or ilomastat (MMPi). **c,** Collagen gel compaction after 24 hours of treatment with either vehicle control or ilomastat. ns = nonsignificant by 2-way ANOVA with Holm-Sidak’s multiple comparisons test.

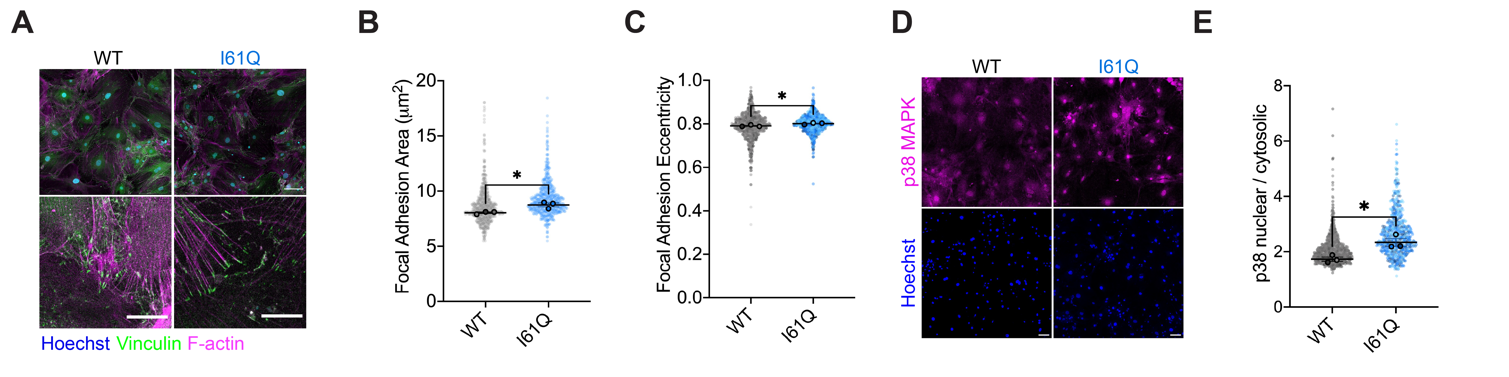

**Fig. S7. Altered focal adhesion morphology and p38 activity are central to the I61Q fibroblast phenotype (A)** Representative images (scale 100μm, inset scale 30μm) of WT and I61Q cardiac fibroblasts stained for vinculin and filamentous actin (F-actin), quantified for **(B)** focal adhesion area **(C)** focal adhesion eccentricity in I61Q fibroblasts and WT controls. **(D)** Representative images from fibroblast immunostaining for p38 MAPK (scale 50μm) and **(E)** quantification for p38 nuclear localization in I61Q and WT fibroblasts, indicating p38 activity. *p<0.05 by unpaired t-test on n=3 biological replicates.

|  | Forward Primer | Reverse Primer |
| --- | --- | --- |
| 18s | GTAACCCGTTGAACCCCATT | CCATCCAATCGGTAGTAGCG |
| Acta2 | ACTGGGACGACATGGAAAAG | GTTCAGTGGTGCCTCTGTCA |
| Ccnd1 | TGCTGCAAATGGAACTGCTTCTGG | TACCATGGAGGGTGGGTTGGAAAT |
| Cdk1 | GGCGAGTTCTTCACAGAGACTTG | CCCTATACTCCAGATGTCAACCGG |

**Table S1.** Primer Sequences used for RT-PCR in this manuscript.

**Data S1. (separate file)**

Normalized RNASeq counts and differential expression results from I61Q and WT cardiac fibroblasts.
